# Uncovering the signaling landscape controlling breast cancer cell migration identifies splicing factor PRPF4B as a metastasis driver

**DOI:** 10.1101/479568

**Authors:** Michiel Fokkelman, Esmee Koedoot, Vasiliki-Maria Rogkoti, Sylvia E. Le Dévédec, Iris van de Sandt, Hans de Bont, Chantal Pont, Janna E. Klip, Erik A.C. Wiemer, Marcel Smid, Peter Stoilov, John A. Foekens, John W.M. Martens, Bob van de Water

## Abstract

Metastasis is the major cause of death in cancer patients and migration of cancer cells from the primary tumor to distant sites is the prerequisite of metastasis formation. Here we applied an imaging-based RNAi phenotypic cell migration screen using two highly migratory basal breast cancer cell lines (Hs578T and MDA-MB-231) to provide a repository for signaling determinants that functionally drive cancer cell migration. We screened ~4,200 individual target genes covering most cell signaling components and discovered 133 and 113 migratory modulators of Hs578T and MDA-MB-231, respectively, of which 43 genes were common denominators of cell migration. Interaction networks of candidate migratory modulators were in common with networks of different clinical breast cancer prognostic and metastasis classifiers. The splicing factors PRPF4B and BUD31 and the transcription factor BPTF were amplified in human primary breast tumors and the expression was associated with metastasis-free survival. Depletion of PRPF4B, BUD31 and BPTF caused primarily down-regulation of genes involved in focal adhesion and ECM-interaction pathways. PRPF4B was essential for triple negative breast cancer cell migration and critical for breast cancer metastasis formation *in vivo,* making PRPF4B a candidate for further drug development. Our systematic phenotypic screen provides an important repository of candidate metastasis drug targets.

## Introduction

Metastasis is the underlying cause of death for the majority of breast cancer (BC) patients. Cancer cell invasion and dissemination, one of the hallmarks of cancer, is driven by aberrant regulation of cell migration (1–3). Cell migration is involved in different steps of the metastatic cascade, including local invasion, intravasation, extravasation, and colonization of secondary sites (4). Although surgery and radiation therapy are generally effective at the primary site, the development of metastatic disease signals a poor prognosis. Therefore, understanding the fundamental mechanisms of cell migration is critical for our comprehension of disease progression.

Cell migration is a highly integrated multistep process initiated by protrusion of the cell membrane in response to migratory or chemotactic stimuli. Cells display several different migration modes, including amoeboid and mesenchymal (also called lamellipodial) migration, as well as multicellular or collective cell migration (5–7). Recent reports have shown that tumor cells display adaptive switching between migration modes, in response to micro-environmental changes (8) or molecular targeting (9, 10). This switching, also referred to as cellular plasticity, makes the metastatic process difficult to target therapeutically, as tumor cells rapidly adapt to changing environments.

Various mechanisms define tumor cell migration, which include signaling components that drive the reorganization of the cytoskeleton, such as Rho-GTPases and its regulators, cell adhesion molecules part of the family of integrin receptors as well as downstream signaling mediators including focal adhesion kinase (FAK) and integrin linked kinase (ILK) (11–13). Often these molecules are part of gene networks that are co-regulated to promote cell migration, such as growth factor-mediated activation of Smad- and ZEB-family members that ultimately drive epithelial-mesenchymal transition (EMT) thereby facilitating cancer cell dissemination (14–17). Several of these individual factors, including FAK, EGFR, HBEGF, ROCK and RHOC are also essential in cancer metastasis (18–23). Independent functional genomics screens have assessed the role of kinases and adhesion-related molecules in cell migration of lung carcinoma H1299 cells (24) and breast cancer MCF10A cells (25). Also recent *in vivo* RNAi screens have identified several novel modulators of cancer metastasis directly (26–28). However, a systematic analysis on the role of the complete spectrum of individual signaling molecules in tumor cell migration is lacking.

Here, we aimed to unravel the landscape of cell signaling components that functionally drive migration of triple negative breast cancer (TNBC) cells, and are likely responsible for cancer cell dissemination and metastasis formation. The TNBC subtype contains ~15% of all breast cancers and has a poor prognosis (29). A large proportion of TNBC cell lines have a mesenchymal phenotype and a high motility behavior in association with a metastatic spread (30, 31). We performed an imaging-based RNAi-screen to identify genes involved in the regulation of different migratory phenotypes in the two highly motile and mesenchymal TNBC cell lines, Hs578T and MDA-MB-231. Given the diverse migratory behavior of these cells, it allowed a more complete coverage of individual genes and gene networks that define TNBC cell migration. Primary hits were extensively validated by deconvolution screens and additional live cell microscopy experiments. These gene sets were used for network analysis, which revealed enrichment of KEGG pathways in cancer and cell migration, and showed a profound overlap with networks derived from cell migration and breast cancer prognostic signatures. Next generation sequencing revealed that PRPF4B, BUD31 and BPTF depletion caused downregulation of various focal adhesion and extracellular matrix (ECM) interaction components. Importantly, PRPF4B knockdown decreases metastasis formation *in vivo* implying an essential role for this protein in breast cancer metastasis formation.

## Results

### A systematic high throughput signaling RNAi screen for TNBC cell migration

To unravel the critical signaling components that drive TNBC cell migration, we selected two most highly motile TNBC cell lines Hs578T and MDA-MB-231 for microscopy-based RNAi screening. These two cell lines both show a mesenchymal type of migration, yet show differences in lamellipodia organization and protrusive activity. These differences are visible during a PhagoKinetic Track (PKT) assay, where cells are seeded on a fibronectin-coated surface covered with a monolayer of beads and allowed to migrate over time (Fig. 1A, Supplementary movies 1 and 2). We optimized the siRNA transfection and PKT assay for full automation by liquid handling robotics and automated microscopy (Fig. 1B and Suppl. Fig. 1). Multi-parametric image analysis of the migratory tracks allowed quantitative assessment of cell migration behavior. To unravel the signaling landscape that regulates mesenchymal tumor cell migration, we focused our screening effort on the complete set of cell signaling components, covering all kinases, phosphatases, (de)ubiquitinases, transcription factors, G-protein coupled receptors, epigenetic regulators and cell adhesion-related molecules. In total, 4198 individual target genes were evaluated in both Hs578T and MDA-MB-231 in 2 biological independent experiments, in which each experiment measured two PKT assay technical replicate plates (see schematic overview of the entire screen set up in Suppl. Fig. 1). Image analysis of single cell tracks was performed using PhagoTracker (32, 33) and quantitative output data was normalized (robust Z-score) to mock transfected control cells using KNIME. High and low Z-scores of individual parameters already showed the effect of siRNA knockdown on cell motility, i. e. low net area or low axial ratio suggests inhibition of cell migration whereas high axial ratio and high major axis indicated enhanced motility (Fig. 1C/D). Even though the quantification provided eight parameters, all the different migratory phenotypes observed within the data were not fully represented by single parameters. Therefore, migratory phenotypes were classified manually and visualized by principal component analysis (PCA)-based clustering (Fig. 1E/F). A combination of parameters was used to define inhibition of cell migration (i.e. small and small round phenotype), loss of directionality (round phenotype), and enhanced cell migration (long rough and long smooth phenotype). Primary hits were identified by setting thresholds on Z-score for the most dominant parameters within each phenotype. For all phenotypes together, we defined in total 1501 primary hits for Hs578T, and 1306 for MDA-MB-231. For each phenotype, the overlap in hits between Hs578T and MDA-MB-231 was determined (Fig. 1G). These overlapping hits (145 in total) showed similar effects on cell migration upon knockdown in both cell lines, suggesting these were bona-fide cell line-independent drivers of tumor cell migration. Hence we selected these overlapping hits for validation by single siRNA sequences. Additionally, to obtain a larger coverage of genes regulating cell migration that would uncover a more cell type specific migratory behavior, we also selected the top 153 hits in each cell line for validation; only genes that have been defined as druggable were validated (Suppl. Fig. 1 and Suppl. Table 1).

**Figure 1.**
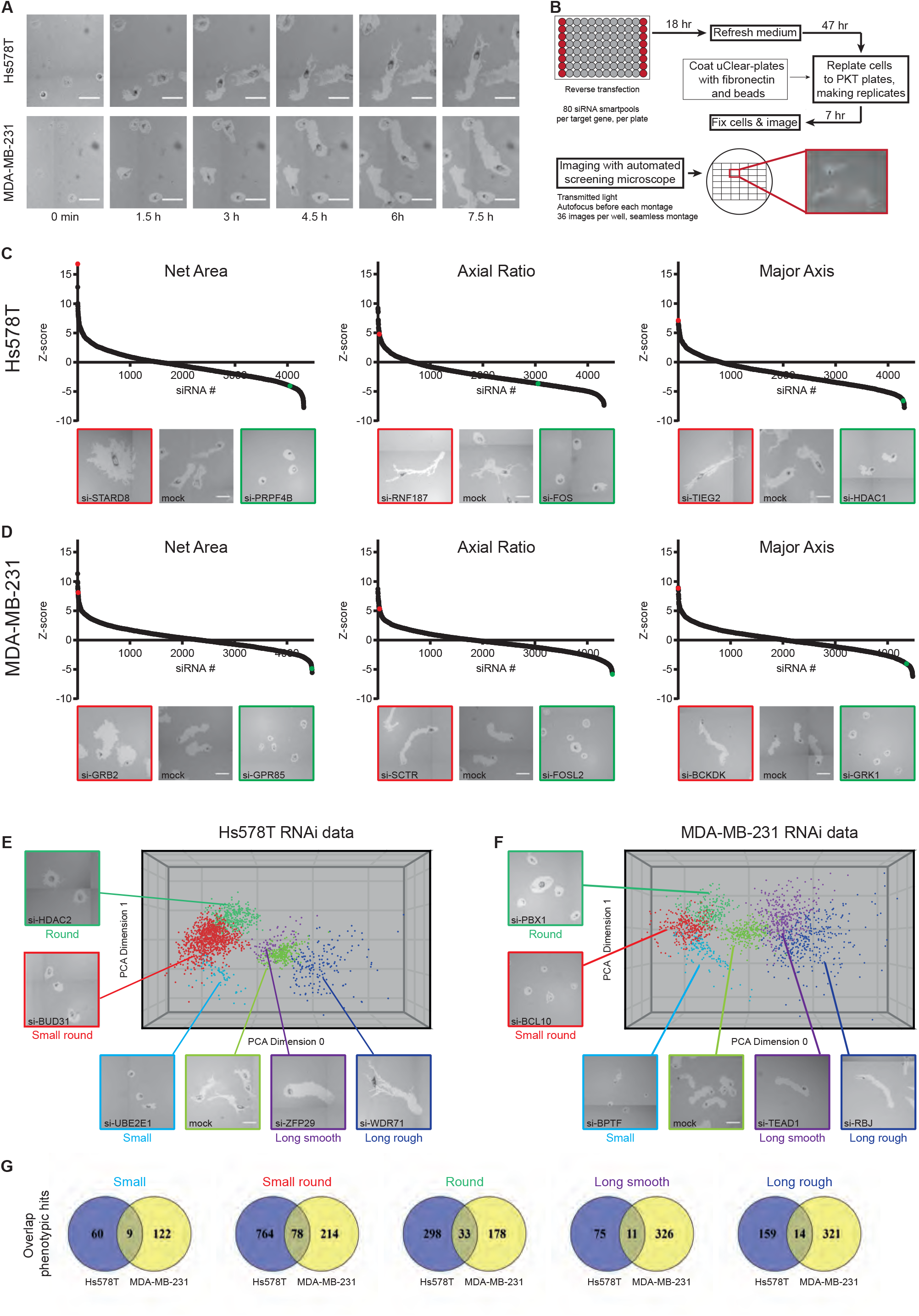
A phenotypic, imaging-based, RNAi screen identifies novel regulators of tumor cell migration. (A) Live cell imaging of Hs578T and MDA-MB-231 phagokinetic tracks. Scale bar is 100 μm. (B) Schematic representation of PKT screen. Transfection was performed in 96-well plates and controls were included in each plate. After 65 h, transfected cells were washed, trypsinized, diluted and seeded onto PKT assay plates. Plates were fixed after 7 h of cell migration and whole-well montages were acquired using transmitted light microscopy. For each siRNA knockdown, a robust Z-score was calculated for each PKT parameter. (C) The three most dominant quantitative PKT parameters are shown for Hs578T and (D) MDA-MB-231. Representative images of migratory tracks for genes with strong effect are shown below each graph and highlighted for enhancement (red) and inhibition (green). (E) Supervised clustering of migratory phenotypes in Hs578T and (F) MDA-MB-231. Migratory phenotypes were identified manually and grouped together in a 3D phenotypic space. Only hits in each phenotypic class and mock control are plotted, and representative images of each phenotype are shown. (G) Overlap of hits in each phenotypic class in both Hs578T and MDA-MB-231.

To validate the primary hits, we repeated the PKT screen assays with both SMARTpool and four single siRNA sequences (Suppl. Fig. 1). First, we confirmed that the SMARTpool knockdown was able to reproduce a similar effect as before and subsequently determined the number of single siRNA sequences showing the same results for the three main phenotypic classes (e.g. small (ZNF446), small round (PRPF4B), and round (PBX1); Fig. 2A). Hits were considered validated when at least 2 out of 4 sequences showed a similar result. Some single siRNAs showed phenotypic switching, i.e. the SMARTpool produced a small round phenotype and the single siRNA showed either a small or round track phenotype; such cases were still considered validated, as these different phenotypic classes indicate a reduction in migration. In total, 217 hits were validated in the Hs578T and 160 in the MDA-MB-231 (for Hs578T see Fig. 2B; for MDA-MB-231 see Suppl. Fig. 2; all validated genes are in Suppl. Table 2). Primary hits that enhanced cell migration in both cell lines were difficult to validate, likely the high motility of our TNBC cell lines is more suited to detect genes that inhibit migration upon knockdown. Indeed, the majority of validated hits were found in the phenotypic classes of reduced cell migration (Fig. 2C). There was an overlap of 65 validated candidate genes that showed inhibition of cell migration in both cell lines (Fig. 2D). Annotation of protein classes for each set of validated hits (Hs578T, MDA-MB-231, and overlap) showed that most of the hits were transcription factors (Figure 2E i) also after correction for library size (Figure 2E ii), suggesting that transcriptional regulated gene networks are critical drivers of TNBC cell migration behavior and consequently metastasis formation.

**Figure 2.**
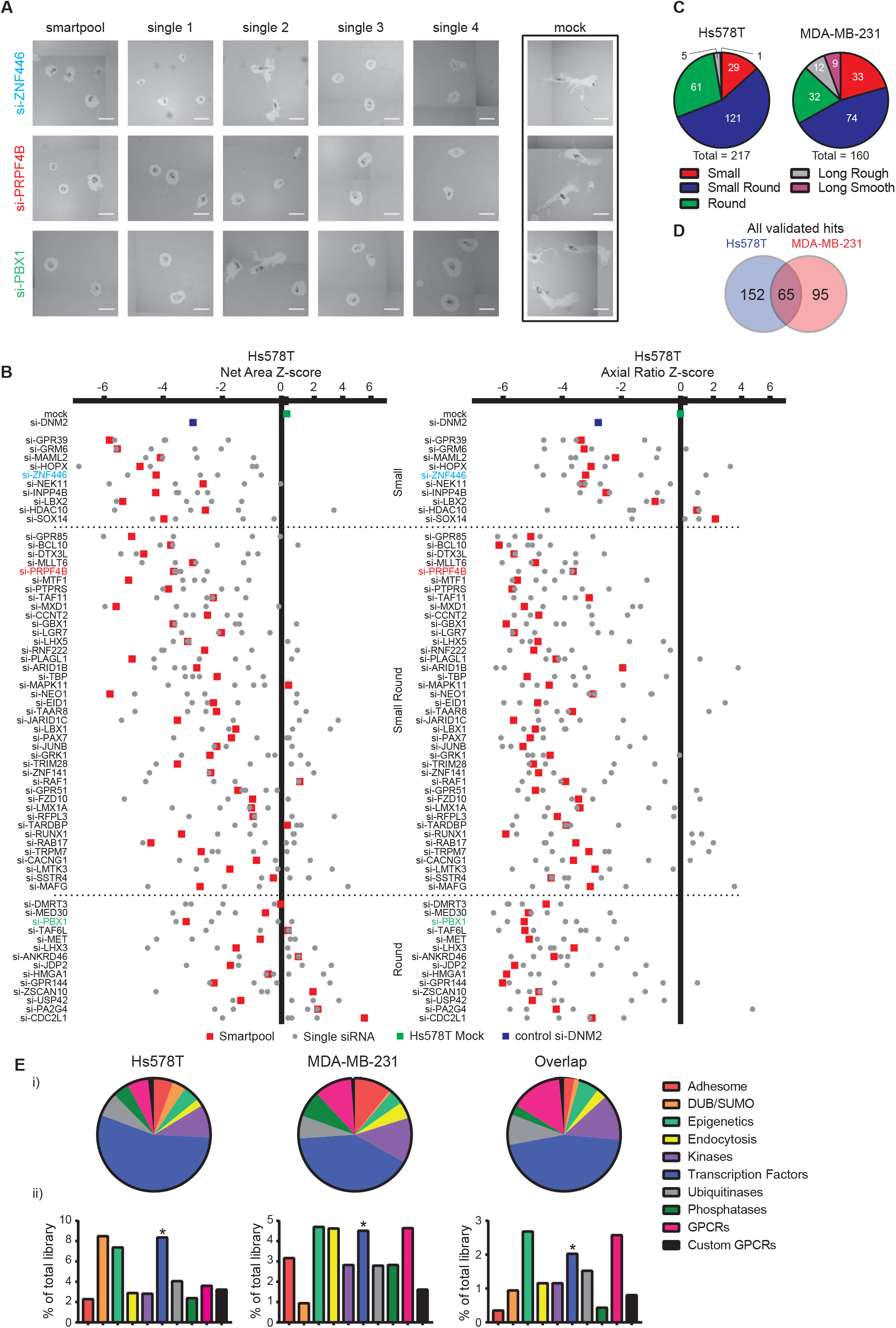
Candidate migratory gene validation by deconvolution PKT screen. (A) Representative images of valid candidate genes using 4 individual single siRNAs in three phenotypic classes are shown. Scale bar is 100 μm. (B) 282 selected genes were tested in a deconvolution PKT screen with 4 single siRNAs per gene. SMARTpool and single siRNA Z-scores of Net Area and Axial Ratio are shown for the ‘overlap hits’ that were validated in Hs578T. (C) Distribution of validated candidate genes in the five phenotypic classes. (D) Overlap of validated hits between Hs578T and MDA-MB-231. (E) Distribution of the validated genes involved in tumor cell migration in Hs578T, MDA-MB-231 over the different libraries based on validated hits effective in both cell lines.

### Transcriptional determinants are critical drivers of BC migratory phenotypes

To further confirm the migratory phenotype detected in the static PKT assay, we evaluated the effect of all validated hits on cell migration using a live microscopy cell migration assay. GFP-expressing Hs578T and MDA-MB-231 cells were used, which allowed fast image acquisition of larger cell populations. Migratory behavior after knockdown was analyzed at the single cell level and normalized to mock (Suppl. Table 3 and 4). Live cell imaging with Hs578T cells confirmed 133 of the 217 hits to inhibit cell migration. Similarly, for the MDA-MB-231 cells, 113 candidate genes (out of 160 validated hits) were confirmed to regulate cell migration. Upon knockdown, 31 PKT overlap candidates inhibited cell migration in this assay in both cell lines (Fig. 3A and 3B and Suppl. Movies 3-14), including various transcriptional and post-transcriptional regulators such as RUNX1, MTF1, PAX7, ZNF141, SOX14, MXD1, ZNF446, TARDBP, TBX5, BPTF, TCF12, TCERG1, ZDHHC13, BRF1, some of which are directly involved with splicing (BUD31 and PRPF4B) or histone modification (HDAC2 and HDAC10). For cell line specific validated hits, we filtered candidate genes for which the expression was associated with clinical breast cancer metastasis-free survival (MFS) in a patient dataset (the Public-344 cohort, GSE5237 and GSE2034, Suppl. Table 5) Many of the hits associated with poor outcome inhibited cell migration in both Hs578T and MDA-MB-231 (Fig. 3A and 3B, see Suppl. Table 3 and 4 for all candidate genes), combined with the overlap candidates resulting in 43 genes that were common denominators of cell migration. Single cell migratory trajectories were plotted for genes affecting cell migration in both cell lines (Fig. 3C). The cell migration movies were visually inspected to determine the effect of knockdown on migration phenotype.

**Figure 3.**
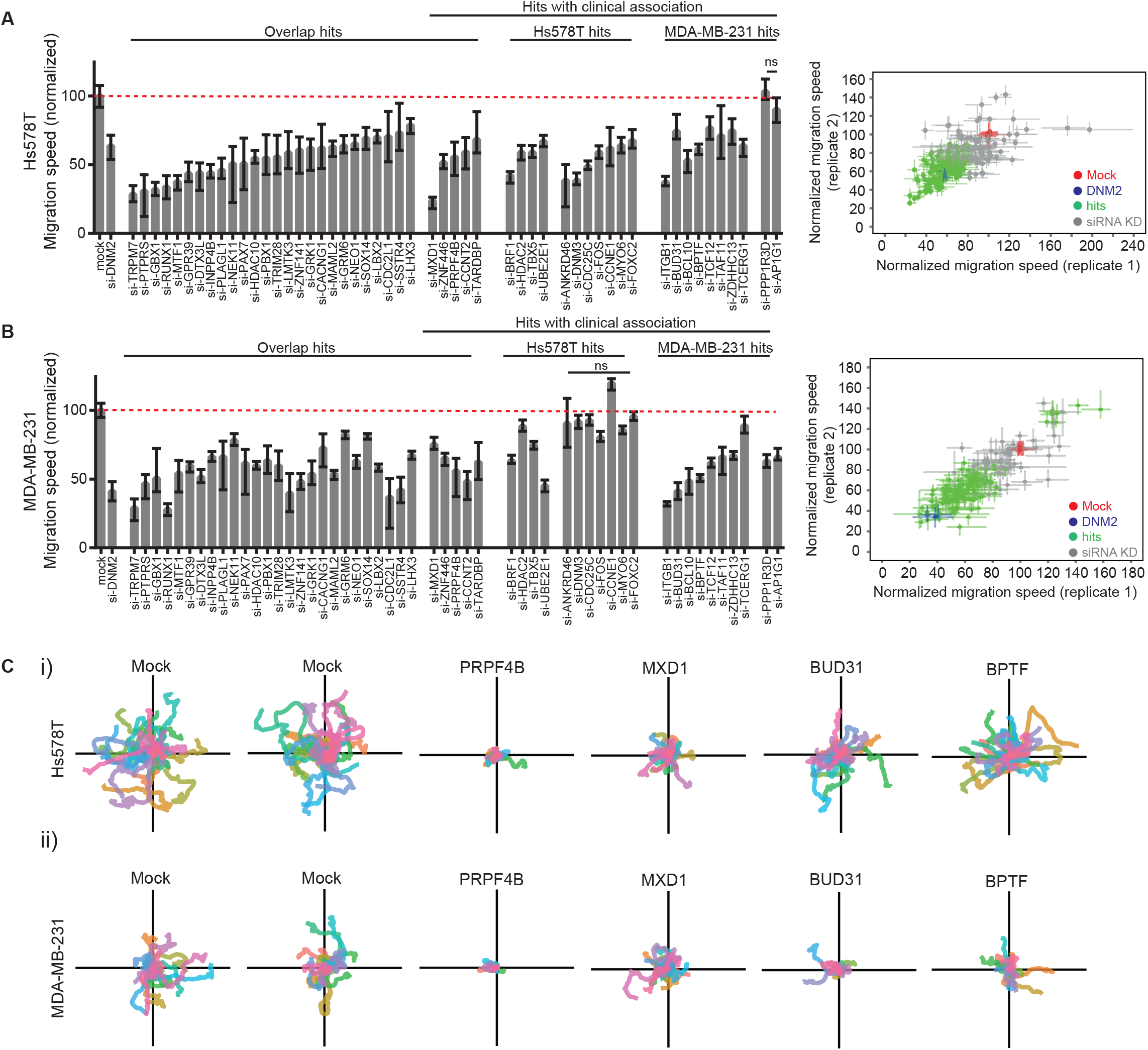
Candidate genes directly affect TNBC cell migratory behavior. (A) Quantification of single cell migration speed of Hs578T-GFP cells after knockdown of validated hits. Hs578T-GFP cells were transfected with siRNAs and cell migration was assessed by live microscopy. Hits were required to show significant and consistent effects in both replicates to be considered as candidate genes (right panel). Median ± 95% confidence interval is shown and cell populations were compared by Kruskal-Wallis test with Dunn’s post correction test. (B) Same as in A, with MDA-MB-231-GFP cells. (C) Single cell trajectories of cell migration upon knockdown of PRPF4B, MXD1, BUD31 or BPTF in (i) Hs578T and (ii) MDA-MB-231 cell lines.

Depletion of PRPF4B and TARDBP resulted in slow and round cells in both cell lines, yet Hs578T cells with knockdown still displayed a dynamic cell membrane with clear protrusive activity (Supplementary movies 5-6 and 15-16). Interestingly, siBUD31 and siUBE2E1 caused very small MDA-MD-231 cell phenotypes that lacked polarization but demonstrated highly dynamic yet unstable protrusions (Supplementary movies 12 and 18); similar effect of BUD31 depletion was also observed in the Hs578T cells. Taken together, our tertiary screen established high confidence in 61% and 71% of our candidate genes (Hs578T and MDA-MB-231 respectively).

### Functional drivers of tumor cell migration partner in networks predictive for BC progression

To better understand the regulatory networks driving BC cell migration, we used the larger lists of our PKT validated candidate genes to inform on protein-protein interaction (PPI) networks that are involved in Hs578T and MDA-MB-231 cell migration. KEGG pathway analysis was performed on the first-order networks of our candidate genes and revealed that similar pathways were affecting cell migration in both cell lines, despite that the networks were constructed from different candidate genes (Figure 4A and 4B, Suppl. Table 6). Several pathways were expected (‘Pathways in cancer’, ‘Focal adhesion’ and ‘Regulation of actin cytoskeleton’) and support that our candidate genes are likely directly connected to signaling networks that determine tumor cell migration. Interestingly, KEGG pathway analysis-based correction of the annotation of the PKT validated candidates not only confirmed the potent role of ‘Transcriptional misregulation in cancer’, but also immune-related, splicing and Notch-signaling pathways in cancer cell migration (Suppl. Fig. 3A). To further investigate the connection of our candidate genes to cell migration and invasion, we correlated our signaling networks to that of three established gene signatures associated with metastatic behavior and cell migration: the Human Invasion Signature (HIS) (34), the Lung Metastasis Signature (LMS) (35, 36), and a 440-gene breast cancer cell migration signature. We previously determined the latter signature based on the gene expression profiles and the migration and invasion capacity of a panel of 52 breast cancer cell lines (manuscript in preparation, Rogkoti et al.). Next, these three independent gene signatures were used to generate PPI networks and the size of each network was reduced to a smaller ‘core’ network (minimum interaction network), which only contained connecting nodes and seed proteins. We next compared the PPI networks from our Hs578T and MDA-MB-231 cell migration screen hits to these three gene signature-based PPI networks (Suppl. Fig. 3B). Both Hs578T and MDA-MB-231 networks show a solid overlap with the 440-gene signature-derived network, with 156 and 145 genes overlapping (Hs578T and MDA-MB-231, respectively). This indicates that the genes we identify as determinants of BC cell migration are part of the gene expression regulons associated with breast cancer cell migration. This notion is further strengthened by the overlap of the Hs578T and MDA-MB-231 PPI networks with the LMS and HIS signature-based networks: 58 (LMS) and 90 (HIS) genes in overlap with Hs578T network, and 53 (LMS) and 77 (HIS) genes with the MDA-MB-231 network. Furthermore, each gene signature derived network showed enrichment for the same KEGG pathways as the PPI networks based on our candidate genes (Suppl. Table 6). Given the high degree of overlap between these three gene signature-based networks and lists of candidate genes, we constructed a single zero-order network based on the combination of candidate genes affecting cell migration in Hs578T and MDA-MB-231 (Fig. 4C). This revealed a sub-network linking 8 transcriptional regulators of which most already have been related to cancer progression, including HDAC2, BPTF, BRF1, TAF11, TCF12 and FOS (37–39).

**Figure 4.**
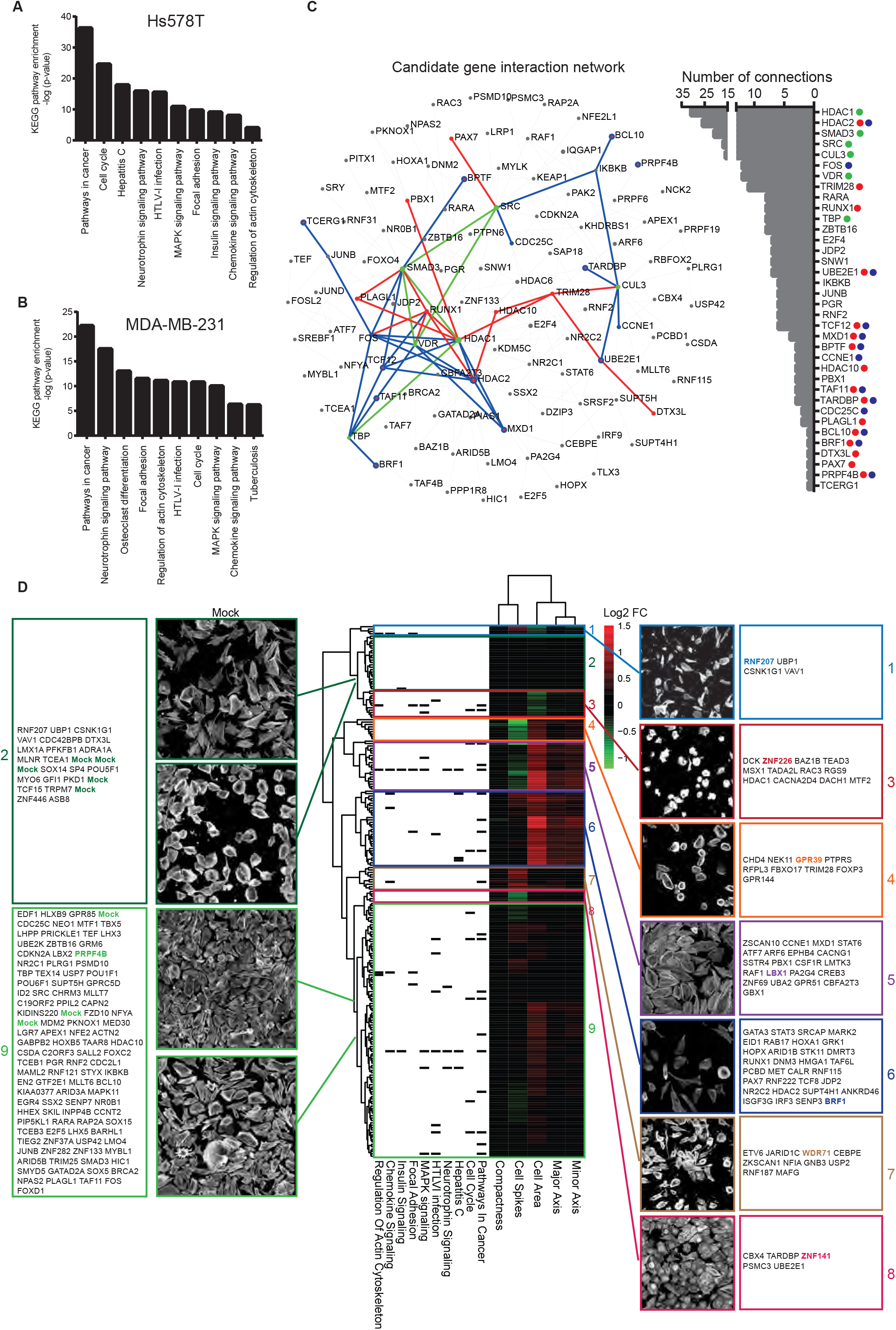
Regulatory networks drive tumor cell migration. (A) Enrichment of KEGG Pathways in PPI networks generated from Hs578T candidate genes and (B) MDA-MB-231 candidate genes. NetworkAnalyst was used to generate PPI networks. (C) Zero-order interaction network of combined Hs578T and MDA-MB-231 candidate genes reveals a highly connected subnetwork of clinically associated genes (in blue). Candidate genes inhibiting cell migration in both cell lines are shown in red; central hubs are highlighted in green. The degree of connectivity (number of connections) is displayed on the right. (D) Phenotype-based clustering of the PKT validated candidate genes based on morphological changes in the Hs578T cell line. Per parameter, log2FC compared to mock control was calculated. Clustering was performed based on Euclidean distance and complete linkage.

Next we systematically investigated the effect of knockdown of the 217 PKT validated hits for morphological changes in the highly polarized Hs578T cell line. 72 hours after knockdown, cells were fixed followed by actin cytoskeleton staining and confocal imaging. For single cells we determined quantitative scores for the minor axis, major axis, cell area, cell spikes and compactness. For each gene knockdown we defined the average scores and fold changes compared to mock control (Suppl. Table 7). Hierarchical clustering grouped our PKT validated hits in nine different clusters (Fig. 4D, see Supplementary Figure 4 for all gene names). Both clusters 2 and 9 contained control knockdown samples but also many genes that affected HS578T cell migration, suggesting that a decrease in migration does not necessarily coincide with an overall change in cell morphology. Not surprisingly, inhibition of cell migration was associated with a wide variety of cellular morphologies: genes in cluster 1 and 3 decreased overall cell area, while genes in cluster 4, 5 and 6 increased overall cell area. Regarding the parameter cell spikes (reflecting the number of cell protrusions), we observed the same variation: some genes reduced cell protrusion formation (cluster 4, 5 and 8), while others enhanced protrusion formation (cluster 7). Our combined data indicate that the various candidate hits differentially affect cell morphology and migratory phenotypes, indicative of different genetic programs that define BC cell migration behavior.

### Modulators of cell migration are associated with BC metastasis-free survival

To further relate our candidate genes to breast cancer progression and metastasis formation in patients, we compared our genes to three prognostic signatures for breast cancer metastasis (40–42). Consistent with previous reports, we observed very few overlapping genes. Despite the minimal overlap of genes, these prognostic gene signatures have many related pathways in common (40). Therefore, we again used the PPI approach to investigate the relation between our migration screen hits and BC progression. Minimum interaction PPI networks were generated based on Wang’s 76 genes, Yu’s 50 genes, and the NKI-70. All three PPI networks showed a robust overlap with our Hs578T and MDA-MB-231 cell migration networks (Fig. 5A and Suppl. Table 8). These three gene expression signatures are strongly predictive of a short time to metastasis, implying that our candidate genes are part of biologically functional regulatory networks and pathways critical in early onset of breast cancer metastasis.

**Figure 5.**
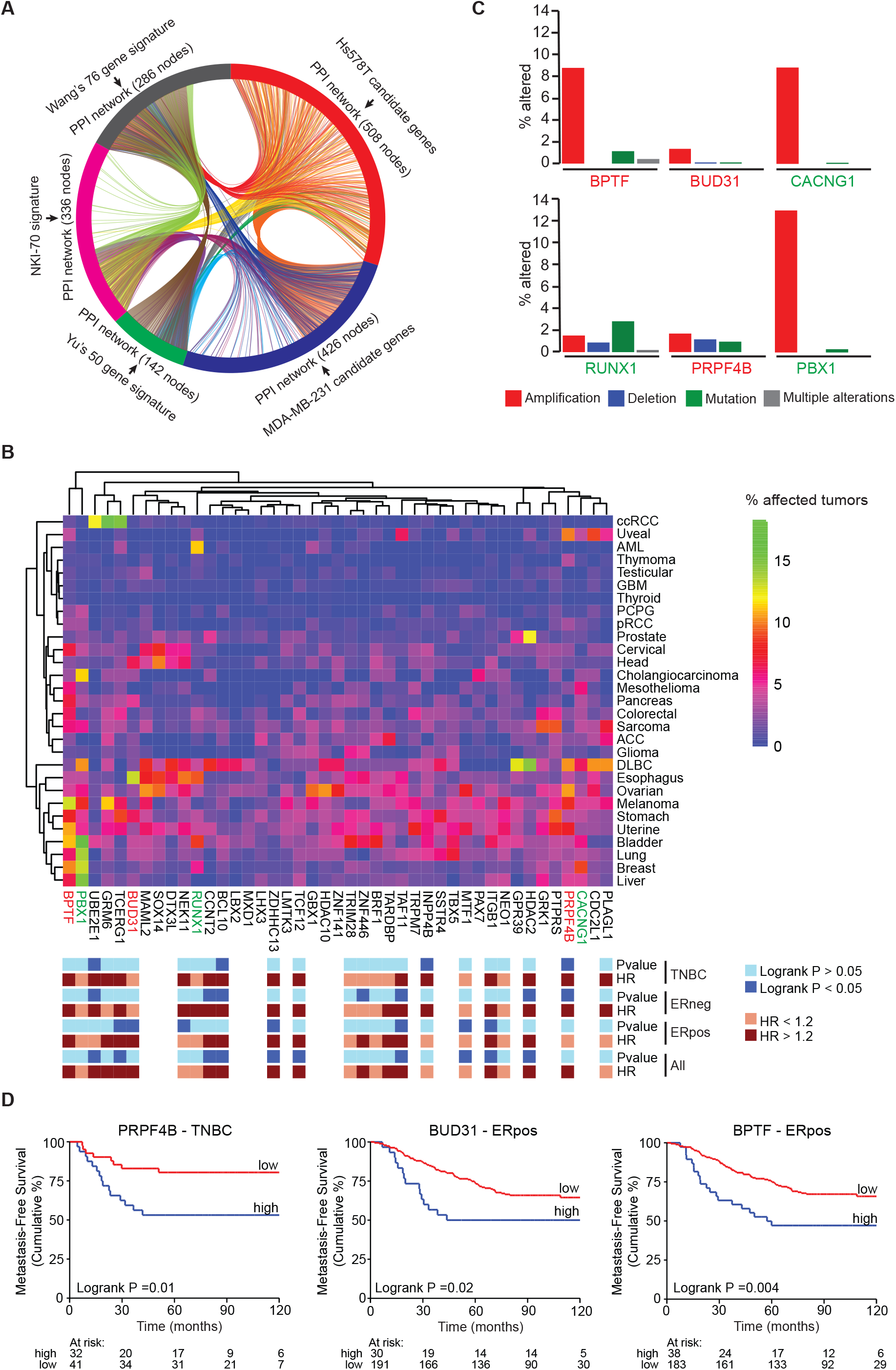
Candidate modulators of TNBC cell migration are related to breast cancer metastasis-free survival and breast cancer progression. (A) Prognostic gene signatures were used to generate minimum interaction PPI networks and compared to our candidate TNBC cell migration gene networks. Candidate genes affecting cell migration feed into similar networks essential for BC progression and metastasis formation. (B) Hierarchical clustering (Euclidean distance, complete linkage) of genetic modifications (mutations, deletions and amplifications combined) of 43 candidate genes in 29 cancer types. Data was derived from The Cancer Genome Atlas. Annotation shows the expression of the candidate in relation to BC metastasis-free survival in different BC subtypes. P-values were calculated using Cox proportional hazards regression analysis, with gene expression values as continuous variable and metastasis-free survival as end point. Genes marked in red and blue are highlighted in C. Red genes were selected for further analysis. (C) Contribution of different genetic modifications to the rate in several highly mutated or amplified candidates. (D) Kaplan Meier curves of for expression of PRPF4B, BUD31 and BPTF and relation to metastasis-free survival in ERpos or TNBC breast cancer. Gene expression data of lymph-node negative BC patient cohort without prior treatment using optimal split was used.

Moreover, we investigated the percentage of mutations, amplifications and deletions (together % altered) of the 43 candidate genes decreasing migration in both cell lines (see Fig. 3A and 3B) in 29 cancer types using publicly available data from The Cancer Genome Atlas (TCGA) (Fig. 5B). We identified clusters of candidate genes highly altered in multiple cancer types, amongst which breast cancer. Separating different types of alterations for the top key factors including BPTF, BUD31, CACNG1, RUNX1, PRPF4B, and PBX1 demonstrated that most of these alterations in breast cancer are dominated by amplifications (Fig. 5C), suggesting that enhanced expression levels of these candidates might be involved in breast cancer initiation or progression. Consequently, we also evaluated the association of the gene expression of these 45 candidate genes with MFS in the Public-344 cohort (Fig. 5D, Suppl. Table 9). 19 out of 24 genes with available expression data showed a hazard ratio >1.2 implicating that high expression of these genes increases the chance of developing metastases with at least 20% in any of the breast cancer subtypes of which 14 are significant (adjusted p-value < 0.05). Interestingly, high expression levels of both splicing factors PRPF4B and BUD31 are associated with earlier metastasis formation in triple-negative and ER-positive tumors respectively (Fig. 5D). Non-core splicing factor PRPF4B is a serine/threonine-protein kinase regulating splicing by phosphorylation of other spliceosomal components (43). PRPF4B has already shown to be involved in taxane resistance and HER2 and ER induced proliferation in BC, but its role in migration and metastasis formation is yet unknown (44, 45). BUD31 is a core splicing factor essential for spliceosome assembly, catalytic activity and associates with multiple spliceosome subcomplexes and has shown to be a MYC target in MYC-driven cancer cells (46). We also identified the transcription factor BPTF, known for its role in chromatin remodeling (47), that is highly amplified in many cancer types and significantly positively correlated to MFS in breast cancer patients (Fig. 5D). BPTF was recently shown to be an important transcription factor interacting with the oncogene MYC in pancreatic cancer (48). We further focused on splicing factors PRPF4B, BUD31 and transcription factor BPTF, since these were newly identified modulators of cell migration associated with BC MFS and/or highly amplified in BC.

### PRPF4B, BUD31 and BPTF modulate expression and activity of cell-matrix adhesion components

In order to identify the mechanisms by which PRPF4B, BUD31 and BPTF were affecting BC cell migration, we performed knockdown of these candidate genes in MDA-MB-231 and Hs578T cells, followed by next generation sequencing (NGS)-based transcriptome analysis (Suppl. Table 10). For all three candidate genes, knockdown efficiency was >90% in Hs578T cells and >80% in MDA-MB-231 cells (Suppl. Fig. 5A). We identified differentially expressed genes (DEGs; log2FC <-1 or >1; adjusted p-value <0.05) for siPRPF4B, BUD31 and BPTF. Notably, expression levels of other validated screen candidates available in our RNA sequencing dataset were not specifically affected by knockdown of PRPF4B, BUD31 or BPTF, indicating that these genes uniquely modulate transcriptional programs that affect cell migration (Suppl. Fig. 6). Knockdown of BUD31 had the broadest effect on gene expression and caused down-regulation of 1119 genes in Hs578T and 929 in MDA-MB-231, with ~50% affected genes overlapping between the two cell lines (Suppl. Fig. 5B-D). PRPFB4 and BPTF knockdown most significantly affected the transcriptome of the Hs578T cells, primarily resulting in down-regulation of gene expression. There was limited overlap in the DEGs between PRPFB4, BUD31 and BPTF (Suppl. Fig. 5E). Since PRPF4B and BUD31 are both splicing factors, we also investigated the effects of knockdown of these candidates on alternative splicing patterns (Suppl. Fig. 7 and Suppl. Table 11). As expected, depletion of BUD31, a core spliceosomal protein, increased the intron retention compared to controls (Suppl. Fig. 7A and 7C) (46). As might be expected from a non-core splicing factor, siPRPF4B only increased a small number of introns retained (Suppl. Fig. 7B and 7C). We were unable to detect any changes in exon inclusion or 3’ or 5’ alterative splice site usage after knockdown of either PRPF4B or BUD31. Since siBUD31-induced intron retention in general relates to reduced expression of the specific gene of interest, we focused on the differentially expressed genes for further analysis. To identify pathways affected by siPRPFB4, siBUD31 and siBPTF we performed KEGG pathway over-representation analysis using the significantly down-regulated genes for all hits in both cell lines separately using ConsensusPathDB (49). Although the overlap in DEGs between different cell lines and knockdown conditions was rather limited, the ECM-receptor interaction was over-represented in all conditions (Fig. 6A, Suppl. Fig. 8A). Moreover, knockdown of PRPF4B, BUD31 and BPTF resulted in down-regulation of the focal adhesion pathway in both cell lines, except for BPTF in Hs578T. Inhibition of these pathways likely directly contributes to the observed decrease in cellular migration. We also observed candidate specific responses: PRPF4B modulates immune signaling (Fig. 6A); BPTF is involved in cell adhesion signaling, as well as general cancer related pathways (Suppl. Fig. 8A); BUD31 regulates a variety of signaling pathways, including metabolic pathways and PI3K/Akt regulated pathways (Suppl. Fig. 8A). Gene set enrichment analysis (GSEA) (50) confirmed the strong down-regulation of the ECM-receptor interaction pathway (Fig. 6B, Suppl. Fig. 8B). Clustering of all genes involved in ECM-receptor interaction (Fig. 6C, see Suppl. Fig. 9 for all gene names) or focal adhesion (Suppl. Fig. 10) demonstrated the involvement of many different pathway components of which some were overlapping between PRPFB4, BUD31 and BPTF (Fig. 6C and 6D); a similar down-regulation was observed at the protein level for several key components in both cell lines (Fig. 6E, Suppl. Fig. 11C). The effects on differential expression of cell matrix adhesion components was also reflected in the different organization of focal adhesions and the F-actin network for both PRPFB4, BUD31 and BPTF (Fig. 6F and Suppl. Figure 11A-B). In summary, both splicing factors PRPF4B and BUD31 as well as the transcription factor BPTF modulate the expression of various focal adhesion-associated proteins and ECM-interaction signaling components in association with distinct cytoskeletal reorganization and decreased BC cell migration.

**Figure 6.**
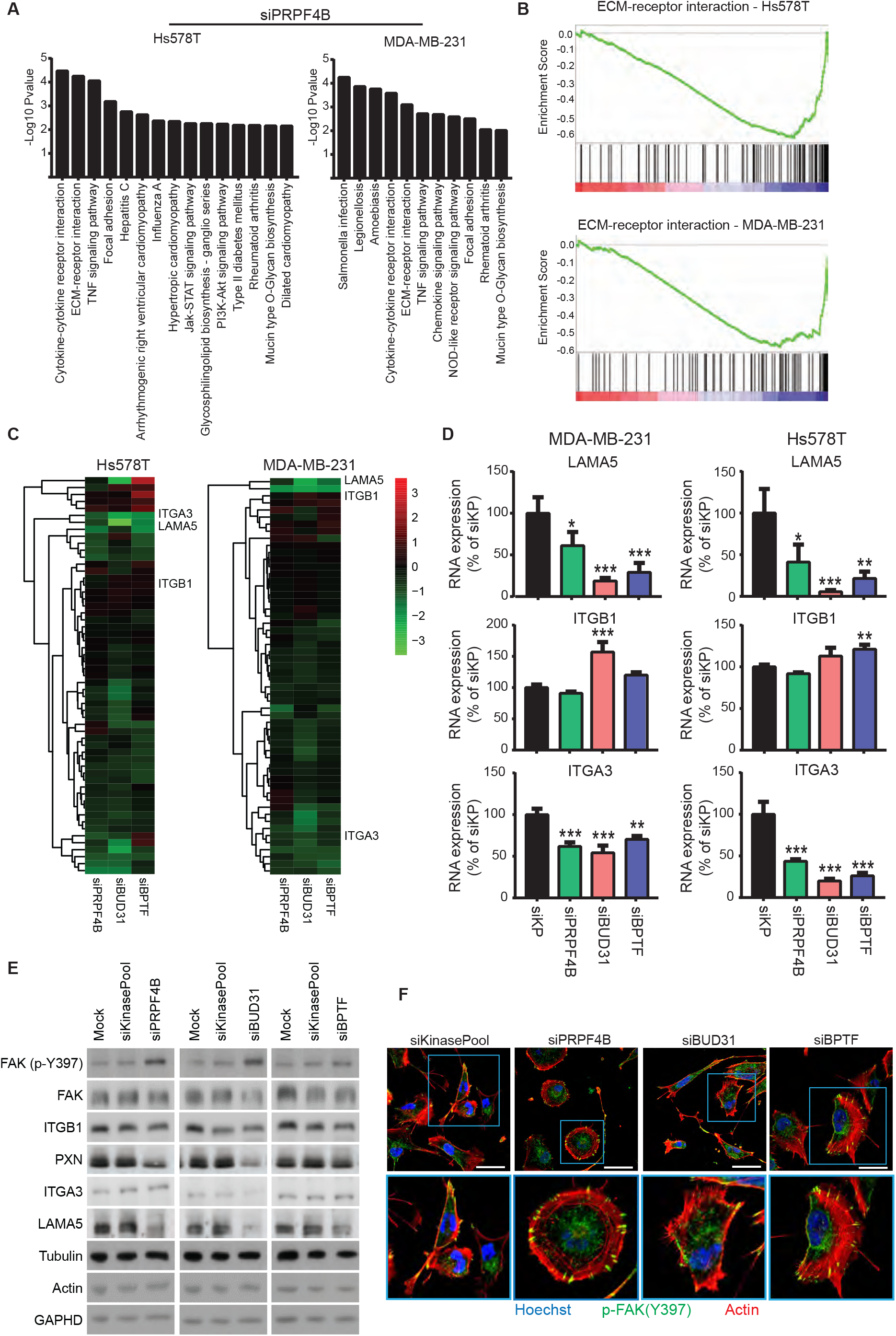
PRPF4B, BUD31 and BPTF depletion affects expression of focal adhesion and ECM-receptor signaling related genes. (A) Over-representation analysis of genes with decreased expression levels (log2FC change < −1) after PRPF4B knockdown in Hs578T and MDA-MB-231 cells using pathways annotated in the KEGG database. (B) Gene Set Enrichment Analysis (GSEA) identifies the ECM-receptor interaction pathway as significantly enriched in downregulated genes after PRPF4B knockdown in Hs578T and MDA-MB-231 cell lines. (C) Hierarchical clustering (Euclidean distance, complete linkage) of log2FC in expression levels of genes involved in the KEGG ECM-receptor interaction pathway after knockdown of candidates demonstrates that many genes involved in this pathway are downregulated after candidate knockdown. (D) LAMA5, ITGB1 and ITGA3 expression upon depletion of PRPF4B, BUD31 and BPTF (one-way ANOVA, *P<0.05, **P<0.01, ***P<0.001). (E) Effect of PRPF4B, BUD31 and BPTF depletion on levels of different ECM and focal adhesion components in MDA-MB-231 cells analyzed with western blot. (F) Effect of indicated gene depletion on focal adhesion and actin cytoskeleton organization. Hs578T cells were fixed and stained against the actin cytoskeleton, p-FAK (Y397). Scale bar is 50 μm.

### PRPF4B is essential for breast cancer metastasis formation *in vivo*

Finally, we investigated whether we could reproduce our *in vitro* findings in an *in vivo* mouse model for BC progression. Using our previously established orthotopic xenograft model, we predicted a decrease in BC metastasis formation upon splicing factor PRPF4B depletion. We selected PRPF4B because its depletion strongly inhibits migration in both Hs578T and MDA-MB-231 cells (see Fig. 3C) and moreover, a role of PRPF4B in metastasis formation has so far not been demonstrated. We established stable PRPF4B knockdown in the metastatic MDA-MB-417.5 cell line that expresses both GFP and luciferase (24, 35, 51). shPRPF4B MDA-MB-417.5 cells demonstrated ~40% PRPF4B knockdown at RNA as well as protein level (Fig. 7A-C). shPRPF4B cells showed an equal primary tumor growth compared to the two shCtrl cell lines (Fig. 7D) which ensured identical time window for tumor cell dissemination from the primary tumor and outgrowth of macro-metastasis (Suppl. Fig. 12A and 12B). Bioluminescence imaging demonstrated that lung metastatic spread was less abundant in the PRPF4B knockdown cells injected group compared to control group (Suppl. Fig. 12C). Both bioluminescent imaging of the lungs *ex vivo* and counting of macro-metastases in the ink injected right lung revealed a significant decrease in metastasis formation in mice engrafted with shPRPF4B cells (Fig. 7F and 7G), which was also confirmed by a decreased lung weight (Suppl. Fig. 12D). *Ex vivo* bioluminescence imaging of the liver, spleen, heart, kidney, uterus and axillar lymph node also showed a decreased metastatic burden by shPRPF4B cells (Fig. 7E) confirmed by a decreased liver and spleen weight (Suppl. Fig. 12E and 12F). Altogether, this demonstrates that PRPF4B knockdown impairs general metastasis formation without showing organ-specificity.

**Figure 7.**
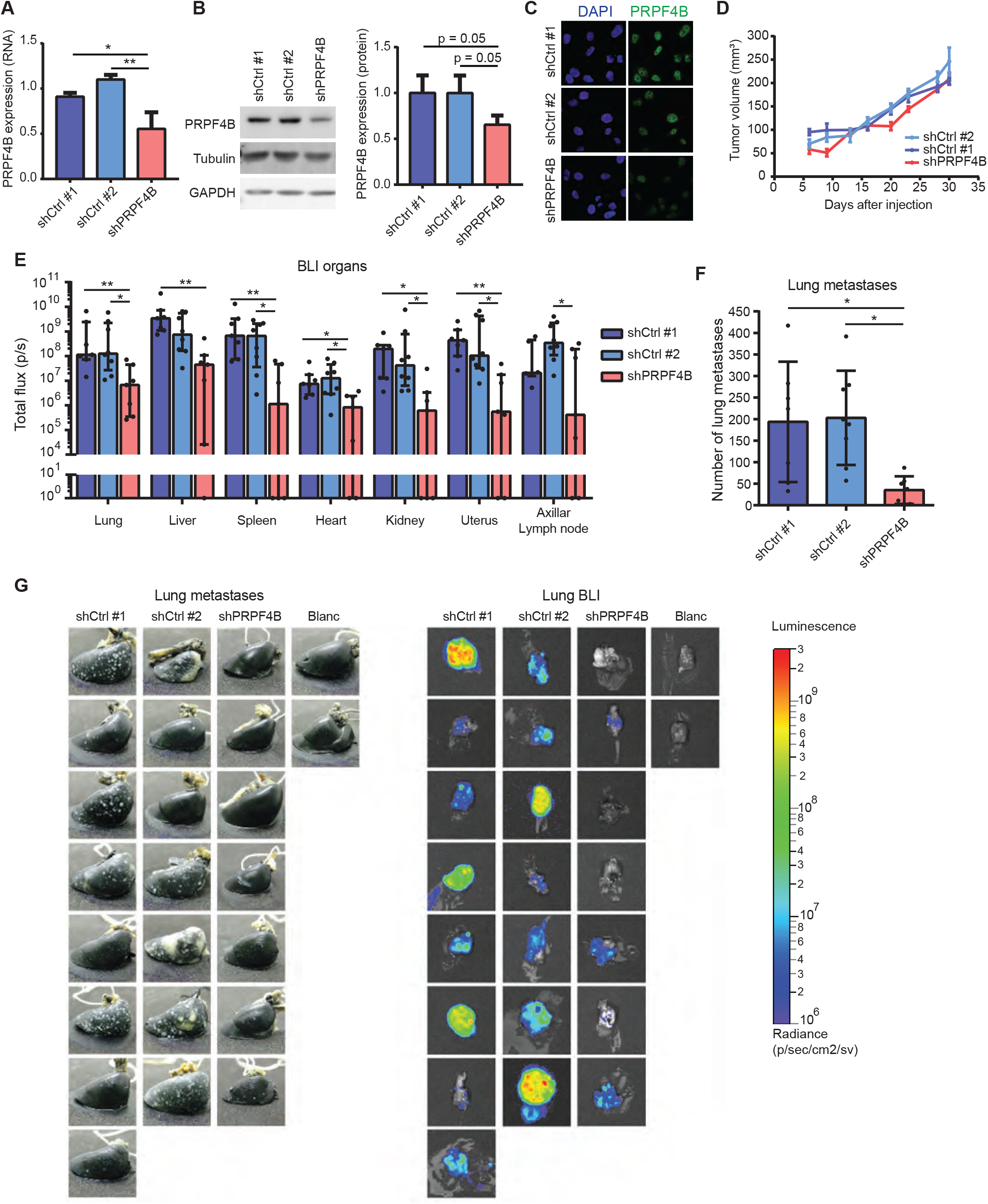
PRPF4B is essential for breast cancer metastasis formation *in vivo.* (A) mRNA and (B) protein expression of PRPF4B in shPRPF4B MDA-MB-417.5 cell line compared to shCtrl#1 and shCrtl#2 MDA-MB-417.5 cell lines (one-way ANOVA, *P<0.05, **P<0.01). (C) Immunofluorescent staining for PRPF4B (green) and DNA (DAPI; blue) in shCtrl#1, shCtrl#2 and shPRPF4B MDA-MB-417.5 cell lines. (D) Tumor growth of shPRPF4B, shCtrl#1 and shCtrl#2 MDA-MB-417.5 cells engrafted in the mammary fat pad of Rag2^-/-^ Il2rg^-/-^ mice (n=7-8 animals per group). (E) Dissemination of shPRPF4B, shCtrl#1 and shCtrl#2 MDA-MB-417.5 cells in the lung, liver, spleen, kidney, heart, uterus and axillar lymph node as determined by *ex vivo* bioluminescent analysis (total flux (p/s)) of different target tissues. (F) Number of lung metastases for shPRPF4B, shCtrl#1 and shCtrl#2 injected mice as determined by macroscopic evaluation of lungs injected with ink. (G) Images of ink injected lungs (left) and bioluminescent signal in the lungs (right) of all individual mice. Blanc indicates mice that did not receive MDA-MD-417.5 cells. Groups were compared by Kruskal-Wallis test with Dunn’s post correction test (luminescence) or ANOVA (metastasis count). * p < 0.05, ** p < 0.01.

## Discussion

New insights into tumor cell migration is highly needed for the identification of potential new drug targets to modulate cancer progression. In the present study, we applied a multi-parametric, high-content, imaging-based RNAi-screen to unravel regulatory networks in tumor cell migration. Our screening effort focused on a broad set of ~4200 cell signaling components, which allowed a high coverage of individual genes and gene networks that affect migration. Our work led to the identification of a defined panel of high-confidence candidate genes that affect the migratory behavior of Hs578T and MDA-MB-231. These genes are part of regulatory networks that are closely related to networks based on gene signatures that are predictive for cell migration, invasion, and metastasis. Our data indicate that different key modulators of cell motility, including PRPF4B, BUD31 and BPTF, define the expression of various genes that compose and regulate cell adhesions. Moreover, PRPF4B, which depletion blocks tumor cell migration *in vitro* is an essential regulator of breast cancer dissemination *in vivo.* Together, our work provides a repository of tumor cell migration drivers and contributes to a broader understanding of the molecular signaling programs that functionally determine BC cell migration and progression of metastatic disease.

Tumor cell migration is a highly heterogeneous and plastic process. Plasticity in cell migration refers to the adaptive and compensatory response to changing environments, which allows tumor cells to switch migration modes when necessary and ultimately metastasize (7). To capture the different migration modes, we performed our RNAi-screen for BC cell migration in two TNBC cell lines. Given the differences in migratory behavior of Hs578T and MDA-MB-231 cells, we anticipated to find novel regulators that affected cell migration independent of migration mode, as well as genes specifically affecting one type of migration. Indeed, the PKT assay allowed us to quantitatively assess different migration phenotypes, as the track morphology reveals the effect on migration, persistence and membrane activity. PCA analysis of the PKT data showed that three migratory phenotypes (small, small round, and round tracks) grouped closely together, resembling different types of inhibition of cell migration. Similarly, two phenotypes of enhanced migration (long rough and long smooth) grouped together on the other side of the PCA space. Interestingly, all these phenotypes were mostly dependent on the parameters net area, axial ratio and major axis, with the enhanced migration phenotypes being separated by track roughness (cell migration with or without high lateral membrane activity). Enhanced cell migration proved to be difficult to validate, with only 8 candidate genes successfully validated and confirmed with live microscopy in the MDA-MB-231 cells. Most of the validated hits decreased net area and/or axial ratio, indicating inhibition of cell migration, as validated by live cell microscopy. The third live microscopy validation resulted in a conclusive selection of validated candidate tumor cell migration modulators: 133 and 113 genes in Hs578T and MDA-MB-231, respectively. The majority of these candidate genes displayed inhibition of cell migration and are most interesting for translation to cancer metastasis. Importantly, candidate migratory regulators, including SRPK1 and TRPM7, have previously been shown to impair cell migration and metastasis formation (24, 52), supporting the robustness of our candidate drug target discovery strategy.

Our work provides a comprehensive resource detailing the role of individual signaling genes in cell migration. Previously, a cell migration screen in H1299 (non-small cell lung carcinoma) identified 30 candidate migration modulating genes (24). Surprisingly, there was little overlap with our validated genes, with the exception of SRPK1. Similarly, little overlap in hits was found with a wound-healing screen in MCF10A cells (25). These differences are likely due to the coverage and size of the screening libraries with the current screen covering ~4200 genes compared to ~1400 genes in the previously published data. Moreover, TNBC cell migration might be driven by different genetic program than non-small cell lung carcinoma and MCF10A cells. Lastly, the MCF10A screen focused on collective cell migration in epithelial cells, which is distinct from single cell mesenchymal migration in our two TNBC cell lines. While our two individual TNBC cell lines showed some overlap in genes that define cell migration, clearly also many distinct molecular determinants were defined for each cell line. This suggest that also the genetic differences between cancer cell lines lead to different molecular networks that can modulate the migratory behavior of tumor cells. For ultimate therapeutic strategies to modulate cancer dissemination the identification of common denominators of migratory and metastatic behavior is essential. Our work contributes to the definition of some of these common players.

The importance of our candidate TNBC cell migration modulators was supported by comparative bioinformatics-based network analysis. While we found little direct overlap between our candidate genes and genes from several different clinical prognostic signatures, our comparative network analyses demonstrated that the cell migration PPI networks in Hs578T and MDA-MB-231 cells are profoundly similar to PPI networks derived from cell migration (Lung Metastasis Signature) (35), cell invasion (Human Invasion Signature) (34), as well as several BC prognostic signatures (40–42). Only cyclin T2, CCNT2, was shared between these signatures and our cell migration data, and was associated with aggressive tumor behavior (40). Moreover, we identified candidates that are highly amplified and/or mutated in many cancer types as well as candidates specifically related to breast cancer metastasis formation. Interestingly, two of these candidates, PRPF4B and BUD31, were splicing factors suggesting modulation of the expression of gene networks through alternative splicing. Moreover, the transcription factor BPTF was one of the major hubs in the interaction network of candidate genes, which is also highly amplified in primary tumors of BC patients and its expression levels are negatively related to MFS. Next generation sequencing revealed that knockdown of PRPF4B, BUD31 and BPTF all resulted in decreased expression of genes involved in focal adhesions and ECM interactions, thus directly driving the observed more rounded and less polarized phenotype in conjunction with decreased cell migration.

Our list of candidate genes that regulate TNBC cell migration likely also contributes to cancer metastasis. We selected PRPF4B to assess whether our *in vitro* screening efforts can be translated to *in vivo* inhibition of metastasis formation. Depletion of PRPF4B almost completely abolished the migration of both Hs578T and MDA-MB-231 cells *in vitro.* Also decreased levels of PRPF4B almost completely eradicated spontaneous metastasis formation from the orthotopic primary tumor to distant organs. This indicates that PRPF4B seems essential in metastasis formation. PRPF4B is a pre-mRNA splicing factor 4 kinase involved in the phosphorylation of PRP6 and PRP31 and splicing complex formation. Yet, in our hands depletion of PRPF4B showed little effect on intron retention patterns, suggesting more subtle splicing effects. In contrast, we found that depletion of BUD31, an important factor in the core of the spliceosome implicated in many catalytic steps of the splicing reaction, drastically increased intron retention in many genes.

Yet, PRPF4B knockdown did affect the expression of various components of the focal adhesion and ECM signaling pathways. Likely, this effect is an important contributor to the reduced migratory and metastatic behavior of TNBC cells. Hence, PRPF4B could be a relevant drug target to combat TNBC dissemination and future research should focus on the development of a specific PRPF4B inhibitor; the x-ray structure of the catalytic domain of PRPF4B suggest this is feasible (53).

Our list of highly confident candidate migratory modulating genes provides ample opportunities for new hypotheses and additional studies in the field of cell migration. 16 G-protein coupled receptors were defined including GPR39, LGR7, GPR85, GRM6 and GPR51. This is particularly relevant as the pathway of GPCR-signaling is one of top over-represented pathways in ER-negative tumors (40). Depletion of several ubiquitinases and proteasome components (DTX3L, UBE2E1, RNF31, RNF115, USP2, USP42, PSMC3, and PSMD10) were also found to inhibit tumor cell migration. Protein homeostasis and proteasome function have recently been suggested as a target for proliferation and growth in basal-like TNBC (54). Transcription factors and regulators make up the largest group of candidate metastasis genes. Some of these factors are part of a large interactive network including HDAC2, BPTF, BRF1, TAF11, TCF12 and FOS. Transcriptional regulation takes place at different levels from direct transcriptional enhancement or suppression, histone modification to pre-mRNA processing thanks to the spliceosome. The downstream targets of these transcription factors are potentially driving BC cell migration, and/or other biological processes that are critical for metastasis formation. Indeed BPTF knockdown affected focal adhesion and ECM components in a similar fashion as PRPF4B and BUD31 knockdown did.

In conclusion, in the present study we used imaging-based phenotypic screening to identify candidate metastatic genes for TNBC that have translational relevance. Understanding the gene networks that are controlled by the various candidate genes provides further insights in the biological programs that define BC cell migration behaviour and lead to important novel drug targets to combat cancer metastasis.

## Materials and Methods

### Cell culture

Hs578T (ATCC-HBT-126) and MDA-MB-231 (ATCC-HBT-26) were purchased from ATCC. MDA-MB-417.5 (MDA-LM2) was kindly provided by Dr. Joan Massagué. All cell lines were grown in RPMI-1640 medium (Gibco, ThermoFisher Scientific, Breda, The Netherlands) supplemented with 10% FBS (GE Healthcare, Landsmeer, The Netherlands), 25 IU/ml penicillin and 25 μg/ml streptomycin (ThermoFisher Scientific) at 37 °C in a humidified 5% CO2 incubator. Stable GFP-expressing Hs578T and MDA-MB-231 cells were generated by lentiviral transduction of pRRL-CMV-GFP and selection of GFP positive clones by FACS. For live cell imaging, phenol red-free culture medium was used.

### Antibodies and reagents

Rabbit anti-PRPF4B (8577, Cell Signaling Technology), mouse anti-Tubulin (T-9026, Sigma-Aldrich), mouse anti-Paxillin (610052, BD Biosciences), mouse anti-ITGB1 (610467, BD Biosciences), mouse anti-FAK (610087, BD Biosciences), mouse anti-N-cadherin (610920, BD Biosciences), mouse anti-E-cadherin (610181, BD Biosciences), rabbit anti-p-FAKY397 (446-24ZG, Thermo Fisher) and mouse anti-Vimentin (ab8069, Abcam) were all commercially purchased. Rabbit anti-ITGA3 and rabbit anti-Laminin5 were kindly provided by A. Sonnenberg (NKI, Amsterdam, The Netherlands). Anti-mouse and anti-rabbit horseradish peroxidase (HRP) conjugated secondary antibodies were purchased from Jackson ImmunoResearch.

### Transient siRNA-mediated gene knockdown

Human siRNA libraries were purchased in siGENOME format from Dharmacon (Dharmacon, Lafayette, CO, USA). Transient siRNA knockdown was achieved by reverse transfection of 50 nM single or SMARTpool siRNA in 2,500-5,000 cells/well in a 96-well plate format (PKT assay and RCM assay, resp.) using the transfection reagent INTERFERin (Polyplus, Illkirch, France) according to the manufacturer’s guidelines. Medium was refreshed after 20 h and transfected cells were used for various assays between 65 to 72 h after transfection. Mock and/or siKinasePool were used as negative control.

### Stable shRNA-mediated gene knockdown

MDA-MB-417.5 cells were transduced with lentiviral shRNA constructs coding for a nontargeting control sequences shCtrl #1 (SHC002), shCtrl #2 (SHC202V) or a sequence targeting the coding region of PRPF4B (target sequence: GCTTCACATGTTGCGGATAAT (TRCN0000426824)) (Mission/Sigma-Aldrich, Zwijndrecht, The Netherlands). The cells were selected by puromycin (sc-108071, Santa Cruz Biotechnology, Heidelberg, Germany). Knockdown efficiency was verified by RT-qPCR, Western Blot and immunofluorescent staining.

### Phagokinetic track (PKT) assay

PKT assays were performed as described before (32, 33). Briefly, black 96-well μClear plates (Greiner Bio-One, Frickenhausen, Germany) were coated with 10 μg/ml fibronectin (Sigma-Aldrich, Zwijndrecht, The Netherlands) for 1 h at 37°C. Plates were washed twice with PBS, using a HydroFlex platewasher (Tecan, Männedorf, Switzerland). Subsequently, the plates were coated with white carboxylate modified latex beads (400 nm, 3.25·10^9^ particles per well; Life Technologies, Carlsbad, CA, USA) for 1 h at 37°C, after which the plate was washed 7 times with PBS. 65 h after siRNA transfection, transfected cells were washed twice with PBS-EDTA and trypsinized. Cells were resuspended into single cell suspensions, then diluted, and finally seeded at low density (~100 cells/well) in the beads-coated plate. Cells were allowed to migrate for 7 h, after which the cells were fixed for 10 min with 4% formaldehyde and washed twice with PBS. For each transfection, duplicate bead plates were generated (technical replicates); transfection of each siRNA library was also performed in duplicate (independent biological replicate). Procedures for transfection, medium refreshment and PKT assay were optimized for laboratory automation by a liquid-handling robot (BioMek FX, Beckman Coulter).

### PKT imaging and analysis

Migratory tracks were visualized by acquiring whole well montages (6×6 images) on a BD Pathway 855 BioImager (BD Biosciences, Franklin Lakes, NJ, USA) using transmitted light and a 10x objective (0.40 NA). A Twister II robotic microplate handler (Caliper Life Sciences, Hopkinton, MA, USA) was used for automated imaging of multiple plates. Montages were analyzed using WIS PhagoTracker (32). Migratory tracks without cells or with more than 1 cell were excluded during image analysis. Quantitative output of PhagoTracker was further analyzed using KNIME. Wells with <10 accepted tracks were excluded. Next, data was normalized to mock to obtain a robust Z-score for each treatment and each parameter. After normalization, an average Z-score of the 4 replicates was calculated. Knockdowns with <3 images were removed, as well as knockdowns with <60 accepted tracks for Hs578T and <150 accepted tracks for MDA-MB-231. Phenotypic classes were identified manually and Z-score thresholds were used as cut-off to define hits.

### Live single cell migration analysis

Hs578T-GFP and MDA-MB-231-GFP cells were transfected with siRNAs as described above and after 65 h, knockdown cell suspensions were seeded in fibronectin-coated black 96-well glass plates (SensoPlate, Greiner Bio-One, Frickenhausen, Germany). Knockdown and control cells were allowed to adhere for 10 h before imaging started. As positive controls, siRNA targeting the GTPase dynamin 2 (DNM2) was used for reduced cell migration and siGFP was used as transfection control. Live microscopy was performed on a Nikon Eclipse Ti microscope, equipped with a 37°C incubation chamber with CO2 flow, an automated xy-stage, Perfect Focus System and 20x objective (0.75 NA, 1.00 WD). Images were captured using a DS-Qi1MC CCD camera with 2×2 binning (pixel size: 0.64μm) and a 2×2 montage was acquired. Two positions per well were selected and GFP images were acquired every 12 min for a total imaging period of 12 h using NIS software (Nikon, Amsterdam, The Netherlands). After live imaging, plates were fixed for 10 min with 4 % formaldehyde and washed twice with PBS. Cells were stained with Hoechst 33258 (H3569, Thermo Fisher Scientific) and Rhodamin-Phalloidin (Sigma-Aldrich) to visualize nuclei and actin cytoskeleton. Nuclei images were captured on the same microscope, acquiring an 8×8 montage of the whole well using a 10x objective. Image analysis was performed using CellProfiler (Broad Institute) (55). For live cell migration, images were segmented using an in-house developed watershed masked clustering algorithm (56), after which cells were tracked based on overlap between frames. Tracking data was organized and analyzed using in-house developed R-scripts to obtain single cell migration data. Only data originating from cells that were tracked for a minimum of 2 h was used. Single cell migration speeds were plotted using GraphPad Prism 6.0 and changes in migration speed were evaluated by comparing cell populations to at least 2 populations of control cells. Two negative control wells with low and high cell densities, comparable to the knockdown populations, were selected for statistical comparison, and knockdowns were required to be statistically significant compared to both controls.

### Imaging-based phenotypic screen

Hs578T cells were fixed and permeabilized in 1% formaldehyde and 0.1% Triton X-100 in PBS and blocked in 0.5% bovine serum albumin (BSA, A6003, Sigma Aldrich) in PBS. Cells were stained with Hoechst 33258 (H3569, Thermo Fisher Scientific) and Rhodamine Phalloidin (R415, Molecular Probes) and imaged using an Nikon Eclipse TE2000-E inverted confocal microscope (Nikon Instruments, Amsterdam, The Netherlands) using a 20x Plan Apo objective, 408 and 561 nm lasers, integrated Perfect Focus System and automated xy-staging. 2x digital zoom, 2×2 stitching images were captured at 4 positions per well. Nuclei and actin cell body were detected by CellProfiler (Broad Institute) and cell area, major and minor axis length, solidity and form factor were measured using the measure object size and shape module in CellProfiler. Images with more than 150 cells were filtered out. Using KNIME, nuclei without a clear cell body were rejected and single cell data was normalized to the median of 2 mock control wells per plate. For the heatmap, all features were mock normalized and clustering was performed on complete linkage and Euclidean distance.

### Immunofluorescence

Cells were fixed and permeabilized 72 h after knockdown by incubation with 1% formaldehyde and 0.1% Triton X100 in PBS for 10 minutes, followed by three times washing with 0.5% w/v BSA in PBS. Cells were incubated with the primary antibody in 0.5% w/v BSA in PBS overnight at 4°C, washed trice with 0.5% w/v BSA in PBS and incubated with the corresponding secondary antibodies and 1:10,000 Hoechst 33258 for 1 h at room temperature. After washing once with 0.5% w/v BSA in PBS and twice with PBS, cells were imaged with a Nikon Eclipse Ti microscope and 60x oil objective.

### Western Blotting

Samples were lysed in RIPA lysis buffer (1% w/w deoxycholate, 50 mM Tris (pH 7.5), 0.15 M NaCl, 0.1% sodium dodecyl sulfate (SDS), 1% v/v NP-40, 2 mM EDTA, 1% v/v protease inhibitor cocktail (P8340, Sigma-Aldrich)) 72 hours after transfection or 2 days after plating for stable knockdown cell lines. Proteins were separated by electrophoresis using SDS-PAGE gels, followed by transfer to PVDF membranes (Merck Millipore), blocked in 5% w/v BSA and overnight incubated with the corresponding primary antibody at 4°C. Membranes were incubated with secondary antibody for 1 hour at room temperature, exposed to Pierce ECL western blotting substrate (Thermo Fisher Scientific) and visualized by using the Amersham Imager 600 (GE Healthcare). At least 2 biological replicates were performed per experiment. Tubulin was used as a loading control.

### RT-qPCR

48 hours after plating stable knockdown cell lines, total RNA was extracted using RNeasy plus mini kit (Qiagen) followed by cDNA synthesis using the RevertAid H minus first strand cDNA synthesis kit (Thermo Fisher Scientific) both according to the manufacturer’s protocol. RT-qPCR was performed with the SYBR Green PCR master mix (Thermo Fisher Scientific) on a 7500 Fast Real-Time PCR machine (Applied Biosystems/Thermo Fisher Scientific) by using the following primers: PRPF4B forward: 5’-CCGAGGAGTCAGGAAGTTCA-3’, PRPF4B reverse: 5’-TCTTTTCAGAATTAGCATCTTCCAT-3’; GAPDH forward: 5’-CTGGTAAAGTGGATATTGTTGCCAT-3’, GAPDH reverse: 5’-TGGAATCAT ATT GGAACAT GT AAACC-3’; β-actin forward: 5’-TCAAGATCATTGCTCCTCCTGAG-3’, β-actin reverse: ACATCTGCTGGAAGGTGGACA-3’. Relative gene expression was calculated after correction for GAPDH and β-actin expression using the 2ΔΔCt method.

### Next generation sequencing

RNA was collected 72 hours after knockdown using the RNeasy plus mini kit (Qiagen) according to the manufacturer’s guidelines. DNA libraries were prepared with the TruSeq Stranded mRNA Library Prep Kit. The DNA libraries were sequenced according to the Illumina TruSeq v3 protocol on an Illumina HiSeq2500 sequencer. Paired-end reads were generated of 100 base-pairs in length. Alignment was performed using the HiSat2 aligner (version 2.2.0.4) against the human GRCh38 reference genome. Gene expression was quantified using the HTseq-count software (version 0.6.1) based on the ENSEMBL gene annotation for GRCH38 (release 84). Count data was normalized and log2 fold changes and adjusted P-values were calculated using the DESeq2 package (57). Calculated log2 fold changes were used to perform ranked gene set enrichment analysis (GSEA) (50). Differentially expressed genes were selected by effect size (log2 fold change bigger than 1 or smaller than −1) and adjusted p-value (smaller than 0.05) and used for over-representation analysis for KEGG pathways using ConsensusPathDB (49).

For the intron retention analysis, RNA-seq reads were mapped to the current human genome (GRCh38) using Hisat 2 (58). Differential intron retention analysis was carried out in R was using DexSeq package (59, 60). In DexSeq the difference of intron inclusion were determined based the counts from the intron and the counts from the two adjacent exons. The sizes of the exons were limited to 100nt immediately adjacent to the intron to reduce artifacts deriving from alternative promoters, alternative splice sites and alternative poly-adenylation sites.

RNA sequencing data is available in Sequence Read Archive with accession number SRP127785.

### Network analysis

Protein annotation of the primary hits was retrieved from QIAGEN’s Ingenuity Pathway Analysis (IPA, QIAGEN Redwood City, USA). Protein-protein interaction (PPI) networks were generated using NetworkAnalyst (www.networkanalyst.ca) (61). Candidate genes were used as seed proteins to construct first-order, minimum interaction and zero-order networks based on the InnateDB Interactome. KEGG pathway analysis was performed on the first-order PPI networks. The connection between multiple PPI networks was visualized by a Chord diagram using NetworkAnalyst.

### Orthotopic mouse model for metastasis assessment

1 x 10^6^ MDA-MB-417.5 shCtrl #1, shCtrl #2 or shPRPF4B cells diluted in 100 μL matrigel (9.2 mg/ml, 354230, batch 4321005, Corning, Amsterdam, The Netherlands) were injected in the fourth mammary fat pad of 7 to 9 week old female Rag2^-/-^ Il2rg^-/-^ mice (n=8 per group). Housing and experiments were performed according to the Dutch guidelines for the care and use of laboratory animals. Sterilized food and water was provided ad libitum and primary breast tumors were surgically removed when they reached the size of 7×7 mm. Next, bioluminescent imaging was used to follow metastasis formation over time. Mice were sacrificed 50 or 51 days after surgery and metastasis formation of all organs was assessed by bioluminescent imaging followed by weighing the lungs, liver and spleen. Finally the right lung was injected with ink in order to count the number of lung macrometastases.

### Breast cancer patient gene expression profiles

Gene expression data of a cohort of 867 lymph node-negative BC patients, who had not received any adjuvant systemic treatment, was used and is available from the Gene Expression Omnibus (accession no. GSE5237, GSE2034, GSE2990, GSE7390 and GSE11121). Clinical characteristics, treatment details and analysis were previously described (40, 41, 62–64). The Public-344 cohort consists of 221 estrogen receptor-positive (ER-positive) and 123 estrogen receptor-negative (ER-negative) patients. Stata (StataCorp) was used to perform Cox proportional hazards regression analysis, with gene expression values as continuous variable and MFS as end point.

### Statistical Analysis

Normality of migration measurements and *in vivo* data was tested using Kolmogorov–Smirnov’s test, d’Agostino and Pearson’s test and Shapiro–Wilk’s test using GraphPad Prism 6.0 (GraphPad Software, San Diego, CA). A data set was considered normal if found as normal by all three tests. Data sets following a normal distribution were compared with Student’s t-test (two-tailed, equal variances) or one-way ANOVA (for comparison of more than 2 groups) using GraphPad Prism 6.0. Data sets that did not follow a normal distribution were compared using Mann–Whitney’s test or a non-parametric ANOVA (Kruskal–Wallis with Dunn’s multiple comparisons post-test) using GraphPad Prism 6.0. Results were considered to be significant if p-value < 0.05.

## Supporting information

## Author contributions

MF, EK, VMR, SELD and BvdW conceived and designed the experiments. MF, EK, VMR, SELD, IvdS, CP and JK performed the experiments. EK and MF analyzed the data. EACW, MS, JAF and JWMM provided the clinical data. PS performed the intron retention analysis. HdB provided technical support for the microscopes. EK and MF wrote the manuscript. BvdW, SELD, JAF and JWMM reviewed and corrected the manuscript.

## Acknowledgements

This project was supported by the EU FP7 Systems Microscopy Network of Excellence (grant no. 258068), the ERC Advanced grant Triple-BC (grant no. 322737) and the Dutch Cancer Society project (grant nr 2011-5124).

## Supplemental data

### Supplemental figures

***Supplemental Figure 1.* RNAi PKT screen setup**. Transfection of up to 10 siRNA library plates per run was performed by automated liquid handling (BioMek). Transfections were performed in duplicate, on different days with separately grown cell cultures. Transfected cells were washed with PBS, trypsinized, diluted and resuspended into single cell suspension, before being seeded in duplicate PKT assay plates (technical replicate). All steps were optimized for automated liquid handling. Whole well montages (6×6) were acquired on a BD Pathway BioImager using transmitted light, and a robotic arm (Twister II, Caliper) placed and removed the PKT assay plates on the microscope. PKT images were analyzed using PhagoTracker software as described previously^20,21^. Quantitative output was normalized to mock control (robust Z-score) using KNIME. Visual inspection of images led to the identification of migratory phenotypes, which were subsequently used for supervised clustering of hits by means of principal component analysis and plotted in a 3D phenotypic space (Fig. 1E,F). Primary hits were selected in two ways: hits that showed overlap between the two cell lines for each migratory phenotype (129 hits) and the top hits affecting cell migration within each cell line (153 hits in Hs578T, 153 hits in MDA-MB-231). Primary hits were validated by deconvolution screens, evaluating the effect of SMARTpool and single siRNA sequences in PKT assays as before. Hits were considered validated if the SMARTpool showed consistent results and at least 2 of 4 single siRNA sequences showed the same phenotype. Ultimately, 217 hits were validated in the Hs578T cells and 160 hits in the MDA-MB-231.

***Supplemental Figure 2.* Phenotypic candidate gene validation by deconvolution screens in MDA-MB-231**. Hits were considered validated if SMARTpool and at least 2 of 4 single siRNAs showed the same effect. 282 selected genes were tested in a deconvolution PKT screen with 4 single siRNAs per gene. SMARTpool and single siRNA Z-scores of Net Area and Axial Ratio are shown for the ‘overlap hits’ that were validated in MDA-MB-231. A list of all the validated hits can be found in the Supplementary Table 2

***Supplemental Figure 3.* Interaction between candidate genes and migration signatures.** (A) Enrichment of KEGG Pathways in Hs578T and MDA-MB-231 validated candidate genes. (B) Chord diagram displaying the connection between our cell migration networks and PPI networks derived from cell migration signatures.

***Supplemental Figure 4.* Phenotype-based clustering of the PKT validated candidate genes based on morphological changes in the Hs578T cell line.** Zoom in of clustering of Fig. 4D. (A) Clustering overview. (B) Zoom in of blue and red clusters shown in A.

***Supplemental Figure 5.* Effect of PRBF4B, BUD31 and BPTF depletion on gene expression**. (A) qRT-PCR of knockdown efficiency of siPRBF4B, siBUD31 and siBPTF used for next generation sequencing in Hs578T and MDA-MB-231 cells. Data are normalized using the ΔΔCT method normalized to actin and tubulin levels. (B) Number of DEGs for siPRBF4B, siBUD31 and siBPTF when compared to control situation (log2FC >1 or <-1) in Hs578T and (C) MDA-MB-231. (D) Overlap of DEGs in Hs578T and MDA-MB-231. (E) Overlap of DEGs comparing different knockdown conditions.

***Supplemental Figure 6.* Effect of PRBF4B, BUD31 and BPTF depletion on expression of other candidate TBNC cell migration modulators**. Hierarchical clustering (Euclidean distance, complete linkage) of log2FC of the gene expression of 35 common validated candidate TNBC cell migration modulators after knockdown of BUD31, PRPF4B or BPTF in Hs578T and MDA-MB-231.

***Supplemental Figure 7.* Effect of BUD31 and PRPF4B depletion on intron retention and exclusion**. (A) Effect of siBUD31 on intron retention. For all introns, the fraction of siBUD31 vs siKP is plotted. Introns for which the inclusion rate significantly differs from siKP are plotted in red (inclusion difference > 0.10 and padj < 0.01). The inclusion difference plotted against the log2FC in expression level demonstrates that intron inclusion in general results in lower expression of these genes. (B) The same as in A, but then for siPRPF4B knockdown. (C) The total number of intron inclusion and exclusion events in siBUD31 and siPRPF4B samples.

***Supplemental Figure 8.* Pathway analysis of altered gene expression after PRPF4B, BUD31 and BPTF depletion**. (A) Significantly over-represented KEGG pathways in the downregulated genes after BUD31 or BPTF knockdown in the Hs578T or MDA-MB-231 cell line. (B) Significantly inhibited KEGG pathways after PRPF4B, BUD31 or BPTF knockdown in Hs578T and MDA-MB-231 cell line identified by performing a ranked gene set enrichment analysis (GSEA).

***Supplemental Figure 9.* Effect of PRPF4B, BUD31 or BPTF depletion on levels of ECM components**. Hierarchical clustering (Euclidean distance, complete linkage) of log2FC in expression levels of genes involved in the KEGG ECM receptor interaction pathway in Hs578T and MDA-MB-231. Zoom in of Fig. 6C.

***Supplemental Figure 10.* Effect of PRPF4B, BUD31 or BPTF depletion of levels of focal adhesion components**. Hierarchical clustering (Euclidean distance, complete linkage) of log2FC in expression levels of genes involved in the KEGG focal adhesion pathway in Hs578T and MDA-MB-231,

***Supplemental Figure 11.* Effect of PRPF4B, BUD31 or BPTF depletion on cell phenotype**. (A) Effect of indicated gene depletion on focal adhesion and actin cytoskeleton organization. Hs578T cells were fixed and stained against the actin cytoskeleton, paxillin (PXN) and Hoechst (nuclei staining). (B) Hs578T cells were fixed and stained against the actin cytoskeleton, p-FAK(Y397) and Hoechst as in Figure 6F. Scale bar is 50 μm. (C) Protein levels of focal adhesion and ECM-interaction related components after knockdown of PRPF4B, BUD31 or BPTF of candidates in Hs578T cells.

***Supplemental Figure 12.* Effect of PRPF4B downregulation on tumor growth and metastasis**. (A) Time of removal of the primary tumor or (B) the total duration of the experiment until sacrifice. (C) Total bioluminescent flux in lungs of mice after surgery till sacrifice. Signal was normalized to lung area. Weight of the (D) lung, (E) liver and (F) spleen after sacrificing the mice. Groups were compared by Kruskal-Wallis test with Dunn’s post correction test (luminescence) or ANOVA (metastasis count). * p < 0.05, ** p <g 0.01.

### Supplemental Movies

***Supplemental movie 1.*** Live cell imaging of MDA-MB-231 (10 frames/second, 4 minutes/frame, 10x objective, 2×2 stitching)

***Supplemental movie 2.*** Live cell imaging of Hs578T (10 frames/second, 4 minutes/frame, 10x objective, 2×2 stitching)

***Supplemental movie 3.*** Live cell imaging of mock control in Hs578T (10 frames/second, 12 minutes/frame, 20x objective, 2×2 stitching).

***Supplemental movie 4.*** Live cell imaging of mock control in MDA-MB-231 (10 frames/second, 12 minutes/frame, 20x objective, 2×2 stitching).

***Supplemental movie 5.*** Live cell imaging of siPRPF4B in Hs578T (10 frames/second, 12 minutes/frame, 20x objective, 2×2 stitching).

***Supplemental movie 6.*** Live cell imaging of siPRPF4B in MDA-MB-231 (10 frames/second, 12 minutes/frame, 20x objective, 2×2 stitching).

***Supplemental movie 7.*** Live cell imaging of siITGB1 in Hs578T (10 frames/second, 12 minutes/frame, 20x objective, 2×2 stitching).

***Supplemental movie 8.*** Live cell imaging of siITGB1 in MDA-MB-231 (10 frames/second, 12 minutes/frame, 20x objective, 2×2 stitching).

***Supplemental movie 9.*** Live cell imaging of siMXD1 in Hs578T (10 frames/second, 12 minutes/frame, 20x objective, 2×2 stitching).

***Supplemental movie 10.*** Live cell imaging of siMXD1 in MDA-MB-231 (10 frames/second, 12 minutes/frame, 20x objective, 2×2 stitching).

***Supplemental movie 11.*** Live cell imaging of siBUD31 in Hs578T (10 frames/second, 12 minutes/frame, 20x objective, 2×2 stitching).

***Supplemental movie 12.*** Live cell imaging of siBUD31 in MDA-MB-231 (10 frames/second, 12 minutes/frame, 20x objective, 2×2 stitching).

***Supplemental movie 13.*** Live cell imaging of siBPTF in Hs578T (10 frames/second, 12 minutes/frame, 20x objective, 2×2 stitching).

***Supplemental movie 14.*** Live cell imaging of siBPTF in MDA-MB-231 (10 frames/second, 12 minutes/frame, 20x objective, 2×2 stitching).

***Supplemental movie 15.*** Live cell imaging of siTARDBP in Hs578T (10 frames/second, 12 minutes/frame, 20x objective, 2×2 stitching).

***Supplemental movie 16.*** Live cell imaging of siTARDBP in MDA-MB-231 (10 frames/second, 12 minutes/frame, 20x objective, 2×2 stitching).

***Supplemental movie 17.*** Live cell imaging of siUBE2E1 in Hs578T (10 frames/second, 12 minutes/frame, 20x objective, 2×2 stitching).

***Supplemental movie 18.*** Live cell imaging of siUBE2E1 in MDA-MB-231 (10 frames/second, 12 minutes/frame, 20x objective, 2×2 stitching).

### Supplemental Tables

***Supplemental Table 1.*** Primary PKT screen. This file contains the full human drugable genome library annotation used for the primary screen, all primary PKT screen data and selected candidate geneIDs and symbols.

***Supplemental Table 2.*** PKT screen validation. Results of the single siRNA validation screen including validated candidates, the resulting phenotypes and number of single siRNAs confirming the smartpool results.

***Supplemental Table 3.*** Live cell imaging in Hs578T. Results of the live cell imaging in Hs578T of PKT derived candidates in Hs578T and MDA-MB-231 candidates that were significantly related to metastasis formation in the Public-344 cohort.

***Supplemental Table 4.*** Live cell imaging in MDA-MB-231. Results of the live cell imaging in MDA-MB-231 of PKT derived candidates in MDA-MB-231 and Hs578T candidates that were significantly related to metastasis formation in the Public-344 cohort.

***Supplemental Table 5.*** Candidate expression related to metastasis formation in human breast cancer patients. This file contains the PKT candidates of which the expression levels were related to metastasis-free survival in ER-negative or ER-postive tumors.

***Supplemental Table 6.*** KEGG pathways in networks. Results of KEGG over-representation analysis of first order PPI networks of the following datasets: candidates PKT screen Hs578T and MDA-MB-231, 440 gene Migration/Invasion signature (440 signature), Human Invastion Signature (HIS), Lung Metastasis Signature (LMS), Yu’s 50 gene signature, NKI-70 signature and Wang’s 76 gene signature.

***Supplemental Table 7.*** Phenotypic screen. Results of phenotypic screen, normalized to mock control.

***Supplemental Table 8.*** PPI network overlap. This file contains lists of genes in the minimum interaction PPI networks established based on candidates PKT screen Hs578T and MDA-MB-231, 440 signature, HIS, LMS, Yu’s 50 gene signature, NKI-70 signature and Wang’s 76 gene signature and the overlap between the different signatures.

***Supplemental Table 9.*** Candidate expression related to metastasis formation in the Public-344 cohort. This file contains all data relating candidate expression levels to metastasis-free survival in human BC patients using the Public-344 dataset. Stata (StataCorp) was used to perform Cox proportional hazards regression analysis, with gene expression values as continuous variable and metastasis-free survival as end point.

***Supplemental Table 10.*** Next generation sequencing of siPRPF4B, siBUD31 and siBPTF in MDA-MB-231 and Hs578T. This file contains basemean (mean count of control and knockdown samples), log2FC and adjusted P-values of all genes for siPRPF4B, siBUD31 and siBPTF compared to siKinasePool control. The DESeq2 package (Love, 2014) was used to normalize the data perform statistics.

***Supplemental Table 11.*** Intron retention of siPRPF4B and siBUD31 in Hs578T and MDA-MB-231. This file contains the intron inclusion differences and adjusted P-values of all introns for siPRPF4B and siBUD31 compared to siKinasePool.

