## Supplementary figures and images for "Uncovering the signaling landscape controlling breast cancer cell migration identifies splicing factor PRPF4B as a metastasis driver"

### Supplementary file 1

#### Supplementary Figure 1

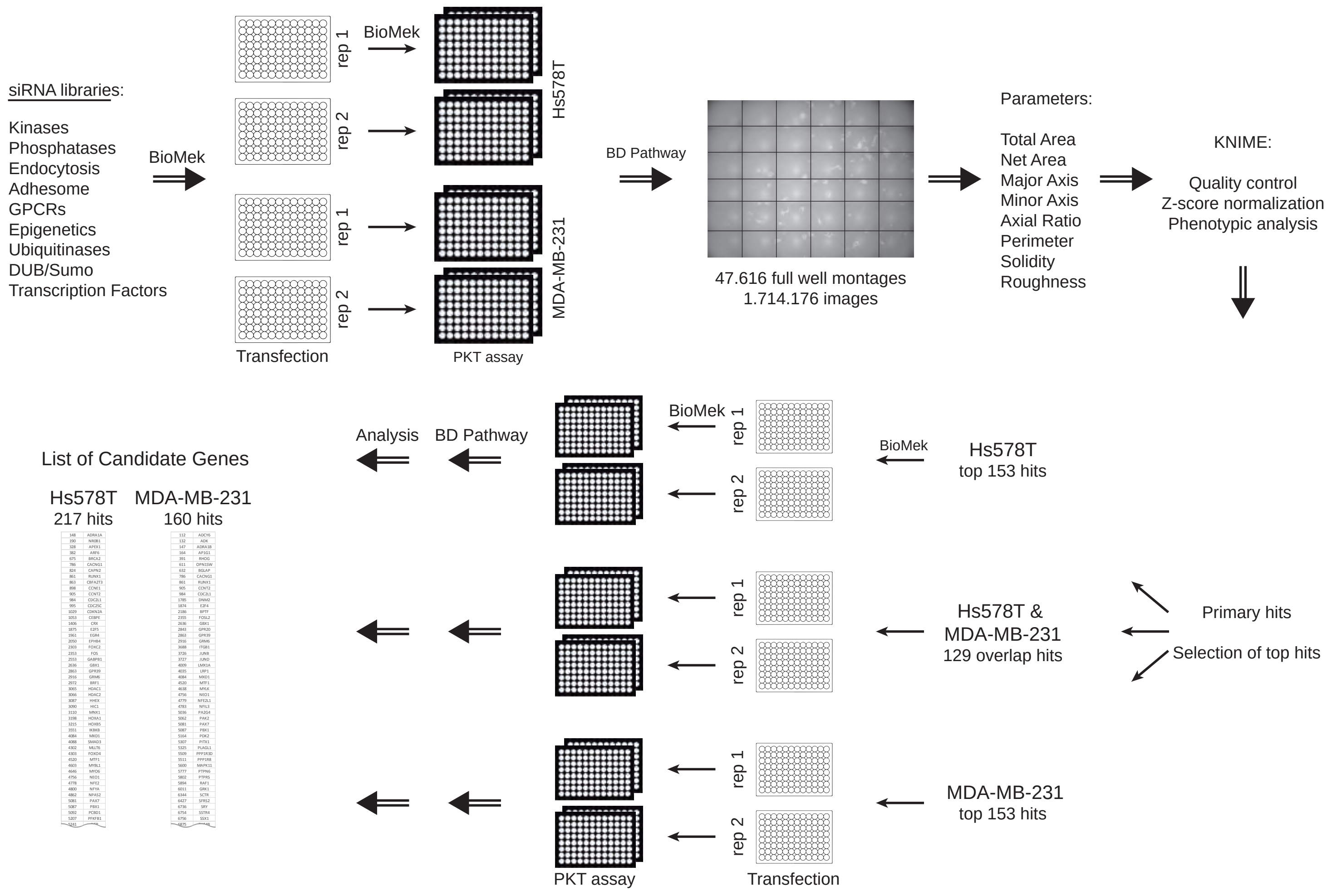

### Supplementary file 2

##### Supplementary Figure 2

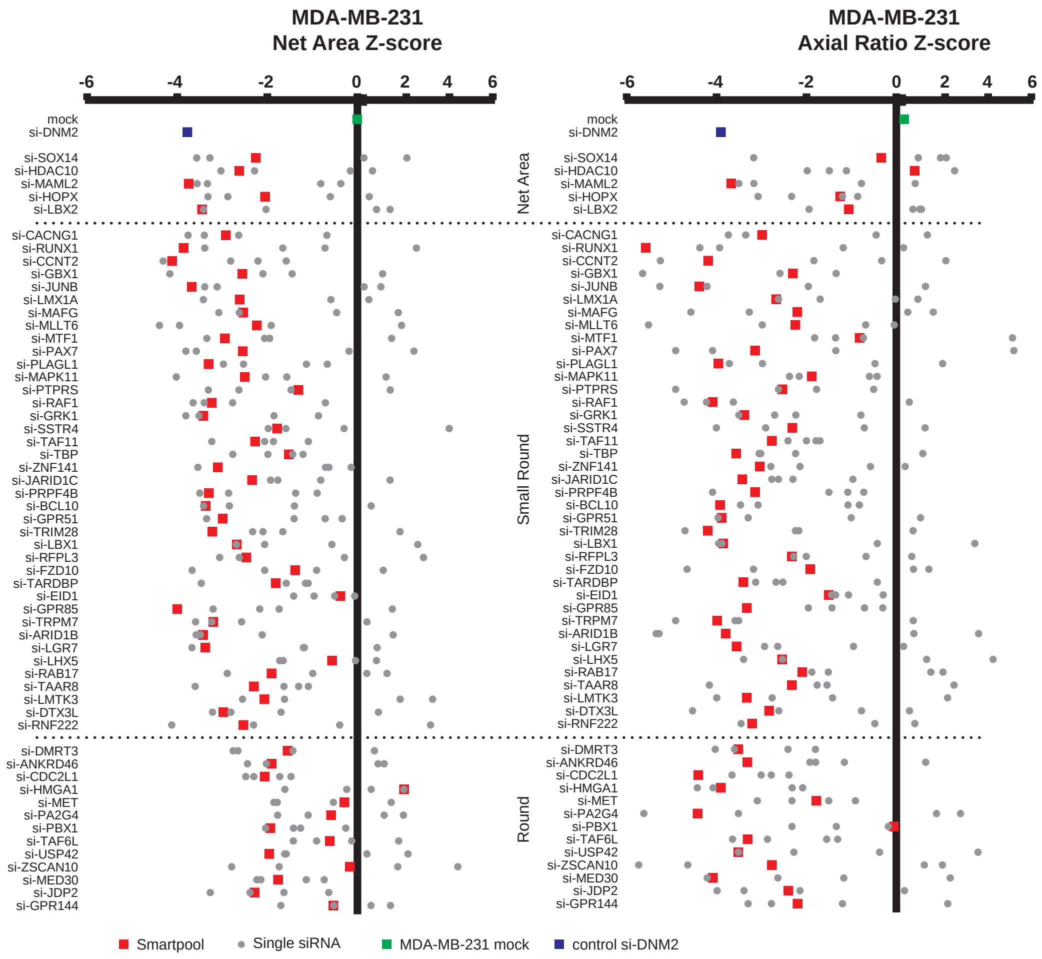

### Supplementary file 3

**Supplementary Figure 3**

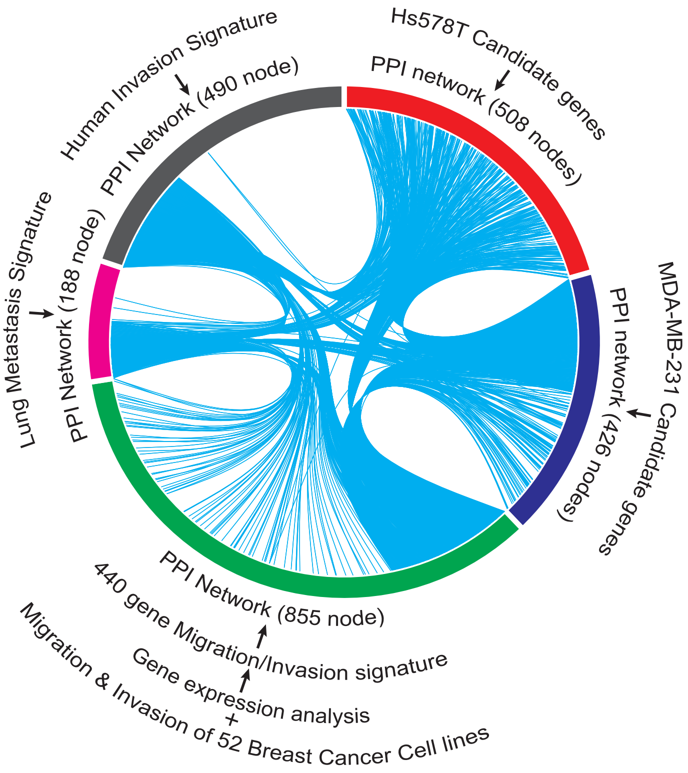

### Supplementary file 4

**Supplementary Figure 4**

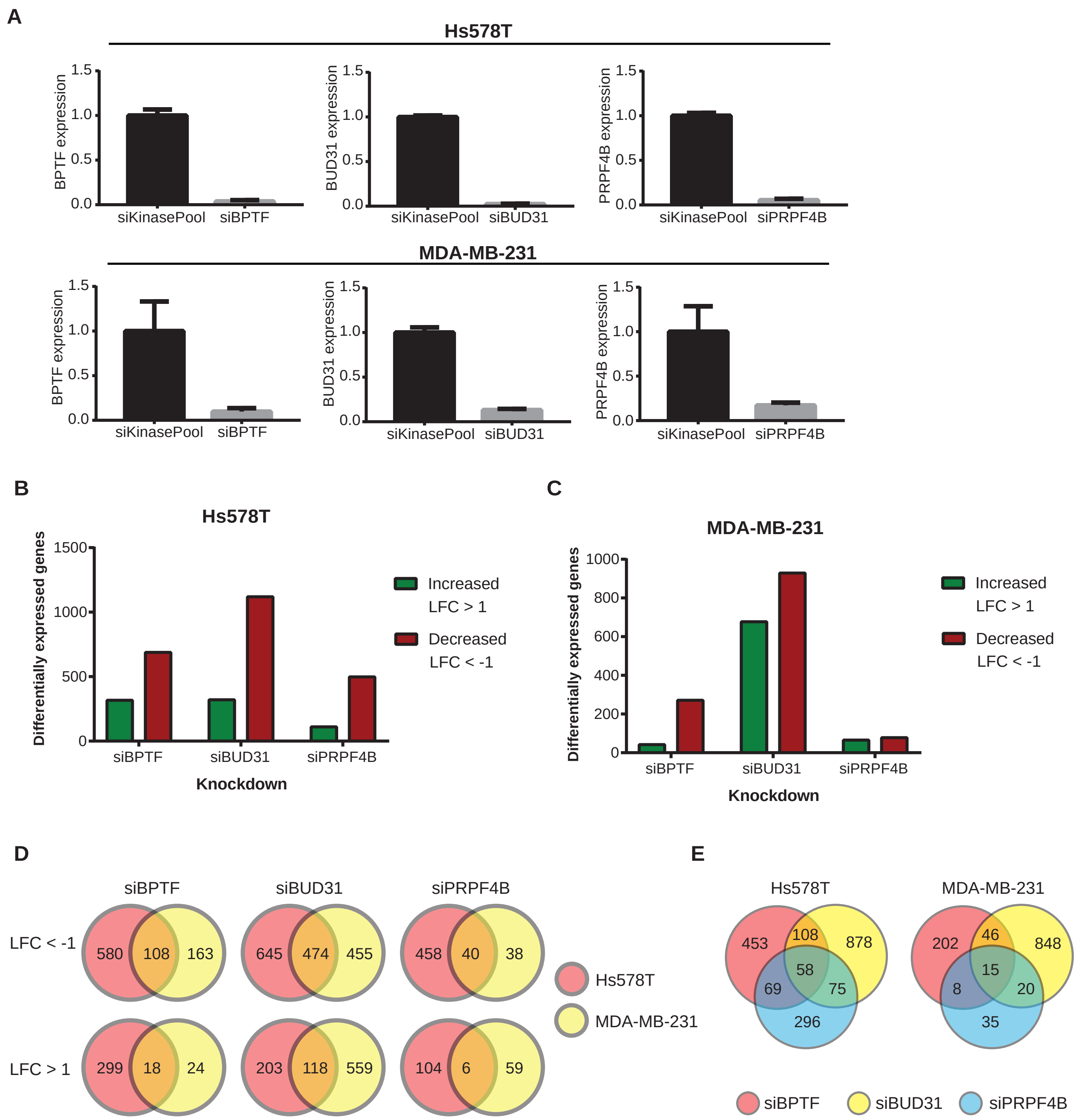

### Supplementary file 5

**Supplementary Figure 5**

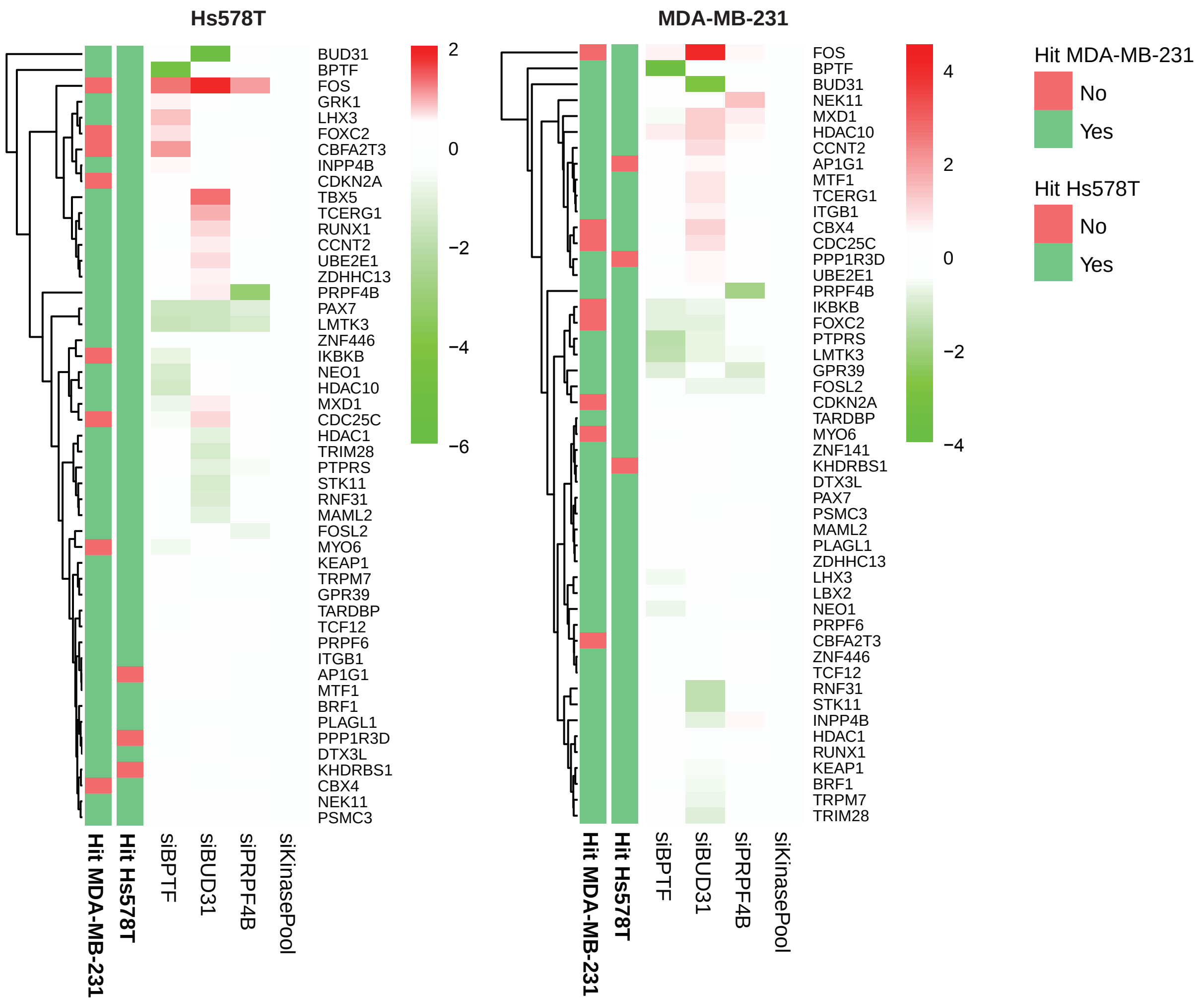

### Supplementary file 6

**Supplementary Figure 6**

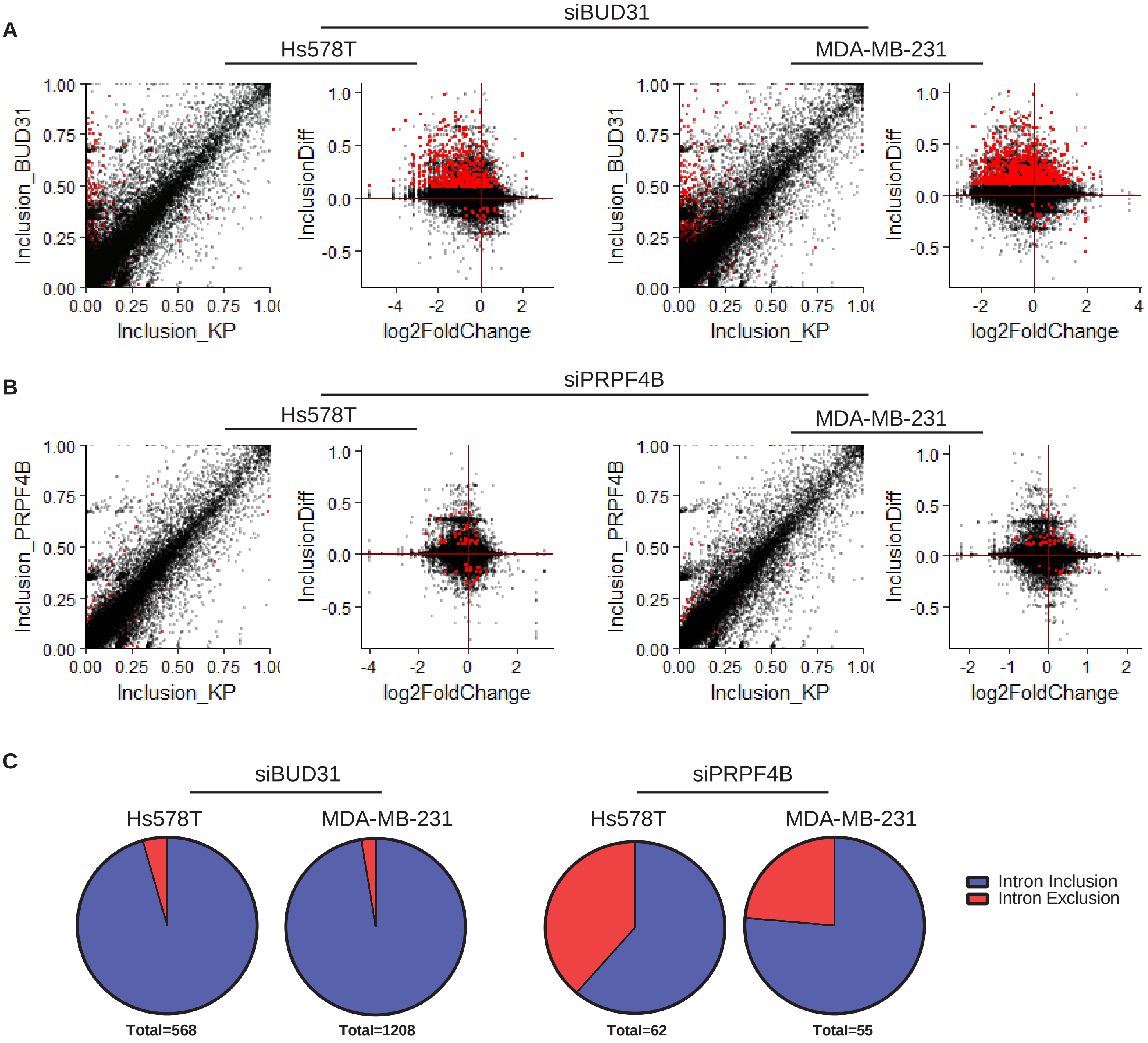

### Supplementary file 7

#### Supplementary Figure 7

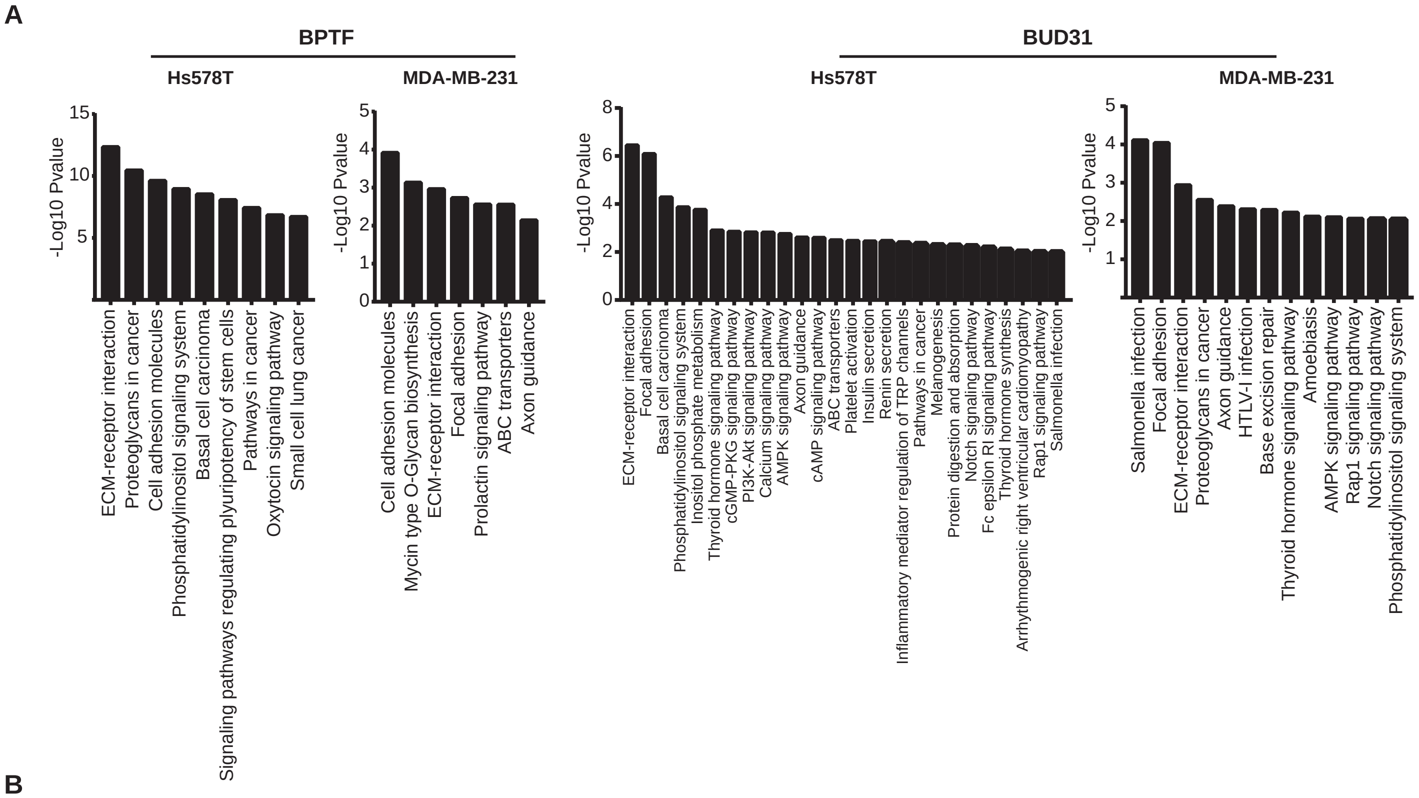


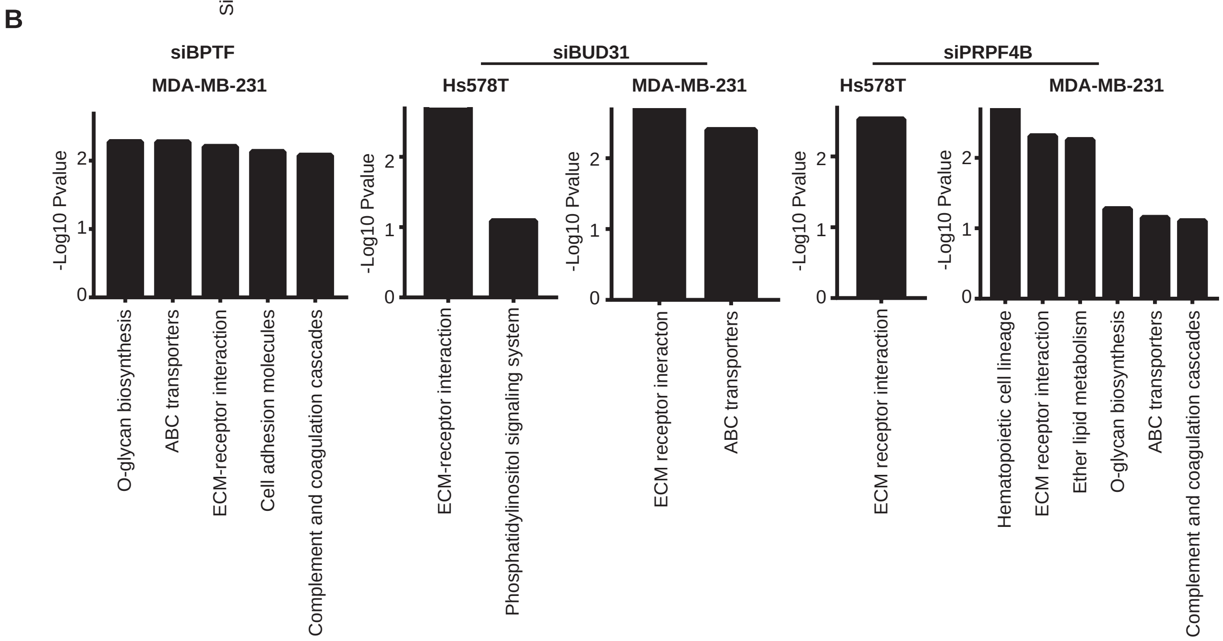

### Supplementary file 8

##### Supplementary Figure 8

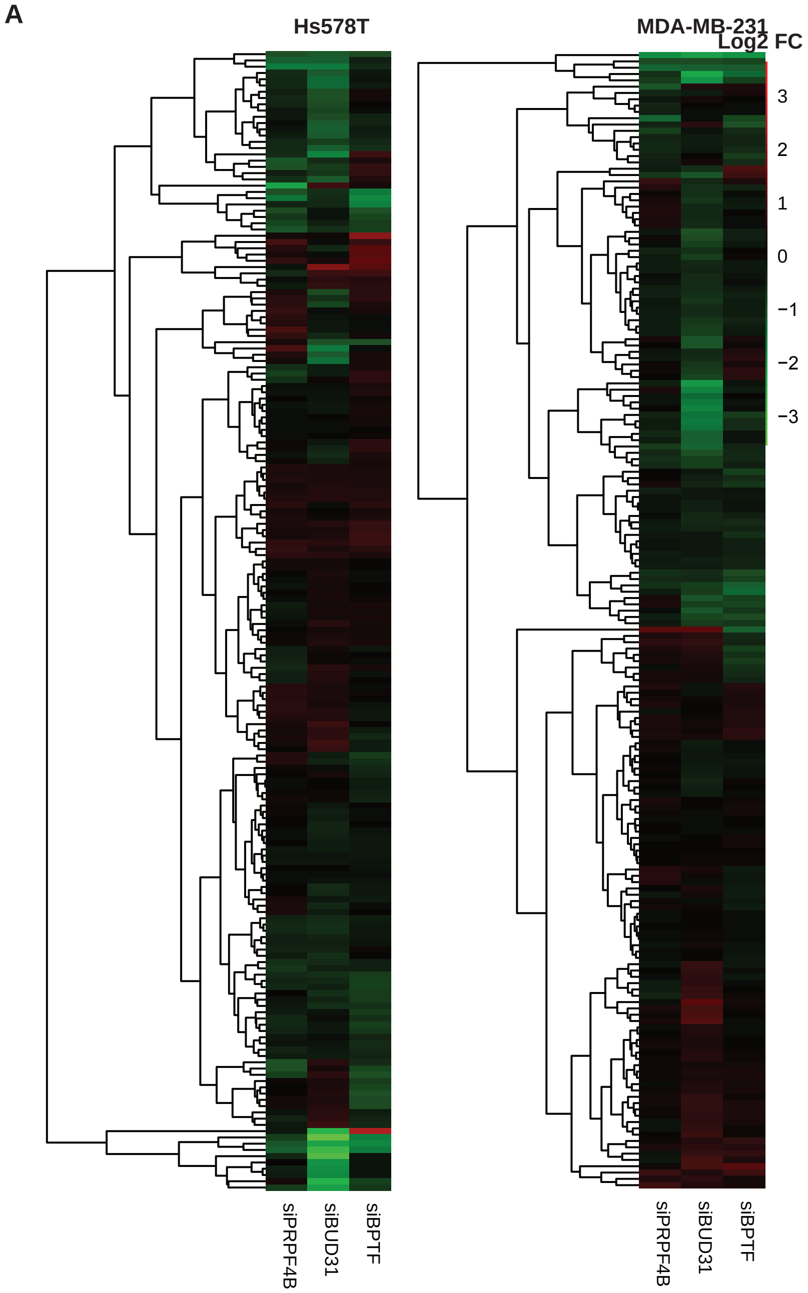


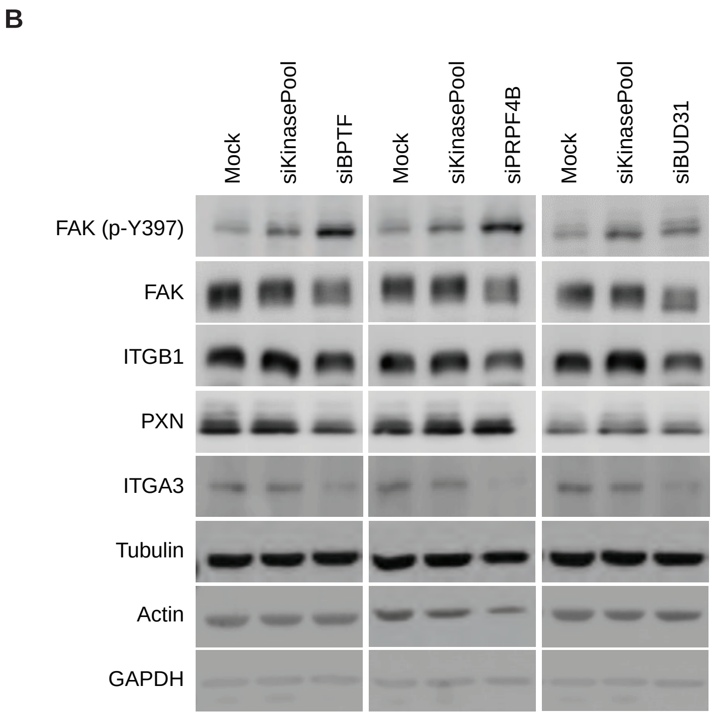

### Supplementary file 9

#### Supplementary Figure 9

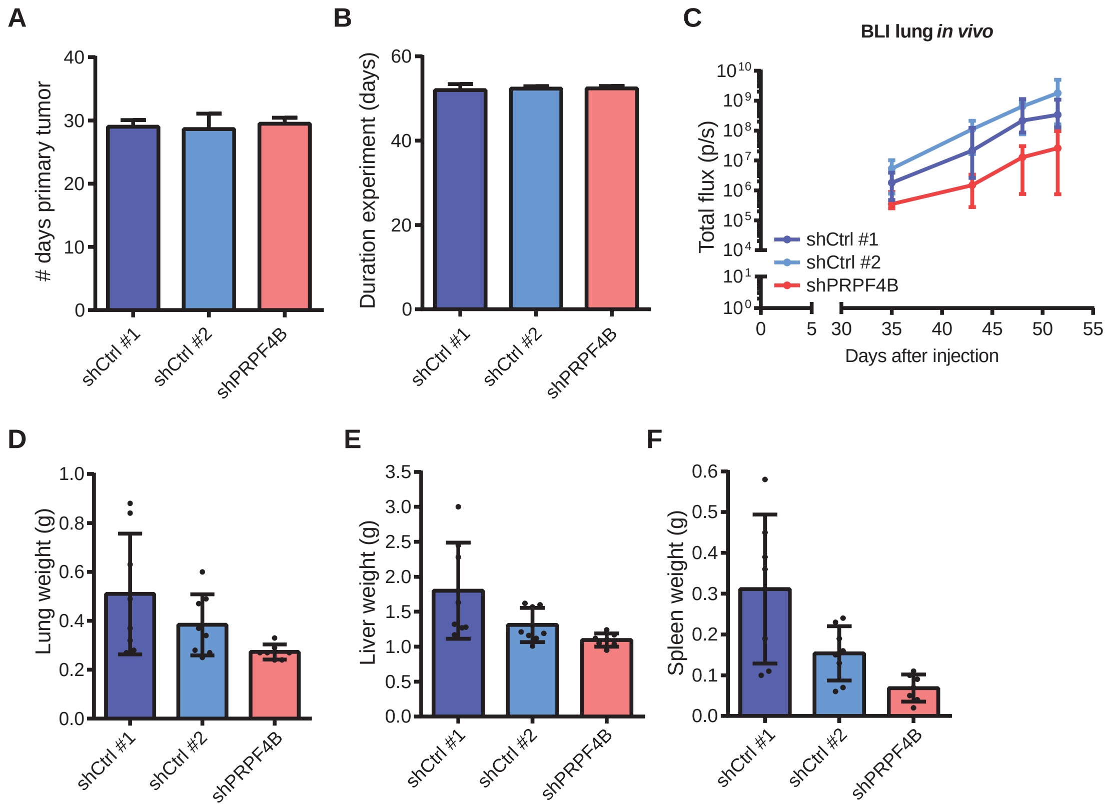
